# Curvature-induced lipid sorting beyond the critical packing parameter

**DOI:** 10.1101/2023.12.15.571845

**Authors:** M. König, R. de Vries, F. Grünewald, S.J. Marrink, W. Pezeshkian

**Affiliations:** Groningen Biomolecular Sciences and Biotechnology Institute and Zernike Institute for Advanced Materials, University of Groningen, Groningen, The Netherlands; Computational Biomolecular Dynamics Group, Max Planck Institute for Multidisciplinary Sciences, Göttingen, Germany; Heidelberg Institute for Theoretical Studies (HITS), Schloss-Wolfsbrunnenweg 35, 69118 Heidelberg, Germany; The Niels Bohr International Academy, Niels Bohr Institute, University of Copenhagen, Copenhagen, Denmark

## Abstract

Over the last few decades, the spatial organization of biomembranes has received a lot of attention. With the fluid mosaic model, Singer and Nicolson proposed that biomembranes behave as two-dimensional fluids, allowing proteins and lipids to diffuse freely and thus react to external stimuli. Biological functionality and membrane organization are intricately linked to each other, however, how these organizations emerge is yet to be established. Here we use coarse-grained molecular dynamics simulations to explore the relationship between membrane shape and lateral organization at the nanoscale for a broad range of lipid mixtures. Our results illustrate that membrane curvature and lateral organization are intimately linked at the nanoscale and the relationship is much more complicated compared to the traditional views, in particular the lipid shape concept. Furthermore, we show that large membrane curvature can disrupt phase separation, line tension, and interleaflet coupling leading to non-intuitive lipid sorting. Our findings highlight how curvature-driven sorting can alter the phase behavior of lipid mixtures, which could be crucial for cellular functionality.

## 1. Introduction

To maintain cellular functionality and ensure intra-cellular homeostasis, biomembranes constantly undergo changes in their chemical composition and physical shape [1]. While long believed to be merely a passive consequence of cellular architecture and function, recent studies have shown that membrane shape is dynamically modulated as an active means to regulate key biological functions [2]. Moreover, a multitude of vital processes involves highly curved membrane intermediates with complex topologies such as endo- and exocytosis, material exchange between the Golgi apparatus and endoplasmic reticulum, and cell fusion and fission [3].

Membrane shape remodeling is regulated by the complex interplay between proteins and lipids at various time and length scales [4, 5]: (i) membrane scaffolding by peripheral membrane proteins with large intrinsic curvature like BAR-domain proteins [6], (ii) pulling forces exerted by the cytoskeleton due to actin polymerization [7] or molecular motors such as kinesins and myosins [8], (iii) insertion of wedge-like amphipathic moieties like the ENTH domain of epsins [9], and (iv) steric pressure arising from macromolecular crowding e.g. of intrinsically disordered domains of endocytic adaptor proteins [10, 11]. The fluid nature of membranes allows proteins and lipids to redistribute and adapt to even extreme changes in membrane shape. While the ability of membrane-bound proteins to generate and sense curvature has been studied extensively [12], the coupling between the spatial distribution of lipids and membrane curvature is yet to be established.

Qualitatively, the lateral segregation of lipids is driven by matching their three-dimensional molecular shape, the so-called intrinsic curvature, and flexibility with the geometrical membrane curvature to minimize the bending energy [13]. For instance, lipids with a small headgroup area relative to their tail cross-sectional area like phosphatidyl-ethanolamine (PE), cholesterol (CHOL), and cardiolipin (CDL), are cone-shaped and have a negative intrinsic curvature. CDL, one of the main lipids in the inner mitochondrial membrane (IMM) of eukaryotic cells [14], segregates into negatively curved (concave) membrane regions as a result of its unique structure consisting of two phospholipids joined at the head. Several experimental studies have shown the localization of CDL to the poles of *E.coli* cells[15-17] as well as the partitioning of CDL to the inner leaflet of DOPC liposomes [18] and to high curvature nanotubes pulled from giant unilamellar vesicles (GUVs) [19].

In addition to the type of headgroup, the degree of tail saturation, which is associated with molecular flexibility and fluidity, also influences the lateral organization. Unsaturated acyl chains with low bending moduli generate a flexible membrane domain with more disordered lipid packing [20, 21], allowing lipids to partition to curved membrane regions and reduce the energy cost of bending the membrane. Mukherjee *et al*. [22] studied the endocytic sorting of fluorescently labeled lipid analogs. They found that narrow endosomal tubes were enriched in short or (poly)unsaturated lipids which can accommodate high degrees of curvature, while long or saturated acyl chains were depleted and showed preferential sorting into more rigid regions.

Despite the simplicity of the model of curvature-induced lipid sorting based on molecular geometry and flexibility, experimental and theoretical work have shown that the energy gain associated with the lateral lipid segregation is not sufficient to compensate for the loss in mixing entropy, resulting in only weak coupling between curvature and lipid sorting [23-26]. The cooperative behavior of lipids is a crucial requirement to overcome the entropic cost of demixing and facilitate curvature-driven lipid sorting. Roux *et al*. [27] reported the formation of a liquid-disordered (*L*_*d*_) domain in membrane nanotubes pulled from GUVs with a homogenous lipid mixture. Following the difference in bending rigidity, the stiffer liquid-ordered (*L*_*o*_) domain, rich in saturated acyl chains, was expelled from the curved nanotube and instead found in the flat membrane region. Further experimental work [28, 29] highlighted the importance of lipid-lipid interactions, concluding that the proximity of the membrane mixture to phase transition is essential for lipid sorting.

During the last two decades, an immense effort has been invested to understand the role of curvature-mediated lipid segregation as one of the organizing principles in biomembranes. However, most experimental studies are based on fluorescence methods that monitor the partitioning of lipid-mimetic dyes in curvature gradients [19, 27, 28, 30, 31]. Such studies can only provide details about the position and local density of the fluorescent dye while the molecular contribution of lipids in the curvature sensing mechanism remains inaccessible.

Molecular simulations have proven to be a powerful tool to overcome this limitation, allowing for the study of complex dynamic processes in biomembranes over a broad range of time and length scales [32, 33]. Thanks to the increase in computational power and efficiency of coarse-grained (CG) molecular dynamics (MD) simulations, studying large-scale membrane shape changes became feasible recently. In line with experimental studies, CG simulations of buckling bilayers, mimicking the membrane composition of the IMM [34, 35] and *E.coli* [36], show the sorting of CDL to highly negatively curved membrane regions. Similar results were obtained from flat bilayer simulations, where negative curvature deflections show strong correlations with CDL enrichment [37, 38]. Despite CDL being the most studied phospholipid in terms of curvature preference, a few computational studies [39-44] predict the sorting of various other types of lipids. For example, the simulation of a large plasma membrane (PM) patch composed of seven different lipid types revealed strong correlations between the negatively curved bilayer regions and clustering of phosphatidylinositol (PI), gangliosides (GM), PE, and CHOL in accordance with their lipid shape [43]. In another study, high membrane curvature is induced by pulling a tether from an asymmetric lipid membrane containing more than 60 different lipid types abundant in the PM [44]. This study showed major differences in the lipid distribution and tether properties depending on the pulling direction. While fully saturated lipids are completely depleted from the tether, (poly)unsaturated lipids were strongly enriched in the curved tether regions independent of the pulling direction and thus independent of the curvature direction.

Similar to the promiscuous behavior of (poly)unsaturated lipids, other factors can drive non-intuitive lipid sorting diverging from the molecular shape concept. For example, CDL can outcompete PE lipids for the most negative curvature at high enough CDL concentrations forcing the PE lipids to occupy regions with zero or positive curvature, despite both lipid types favoring concave bilayer regions [36]. Non-additive curvature preferences for lipid mixtures have also recently been established in a systematic study by Sodt and coworkers [45]. Moreover, fascinating sorting behavior can arise from the interplay between curvature and membrane tension. When compressing a small phase-separated vesicle, the rigid *L*_*o*_ domain partitions to the highly curved region indicating a softening of the *L*_*o*_ phase upon bending [46].

Despite numerous efforts in recent years to elucidate the lateral organizing of curved membranes, it remains still unclear if matching lipid shape with the membrane curvature is sufficient to induce lipid segregation in biological membranes. Here we use CG Martini simulations to systematically characterize lipid sorting and derive a unified relationship between lipid shape and membrane curvature. We show that sorting of CDL in curvature is a direct consequence of its unique shape, however, the degree of sorting strongly depends on various factors like pH, counterions, concentration, and lipid environment. Furthermore, we address the limits of the molecular shape concept and elucidate the non-additive effects that lead to non-intuitive lipid sorting including the effects of lipid packing, lipid phase separation, interleaflet coupling, and line tension.

The rest of this paper is organized as follows: First, we show that our method of confining membranes does not affect lipid organization and dynamics. Then we proceed by studying binary mixtures of PC and CDL lipids and illustrate how CDL’s preference for negative curvature can be modulated. We also comment on the homogenizing effect of lipid mixing entropy and show that curvature-induced sorting of CDL is not hindered by lipid mixing. Next, we expand our study to binary mixtures of various other phospholipids and extract common sorting principles. Finally, we demonstrate that the molecular shape concept is not always sufficient to predict the curvature preference, but that other factors like lipid packing and lipid-lipid interaction can overrule the intrinsic curvature preference.

## 2. Results

### 2.1 The presence of a confining wall does not have a significant impact on the bilayer properties

The desired membrane curvature during the MD simulations is maintained by scaffolding wall particles, placed at the level of the lipid headgroups. These particles interact exclusively with the lipid acyl chain beads in a repulsive manner (see Methods). To assess the influence of the presence of a wall on the membrane structure and dynamics, we set up two flat POPC membranes, with and without (as a control system) confining wall beads. Subsequently, we computed and compared the density profiles, bilayer thickness, area per lipid, and tail order parameter (**Figure 1**).

**Figure 1.**
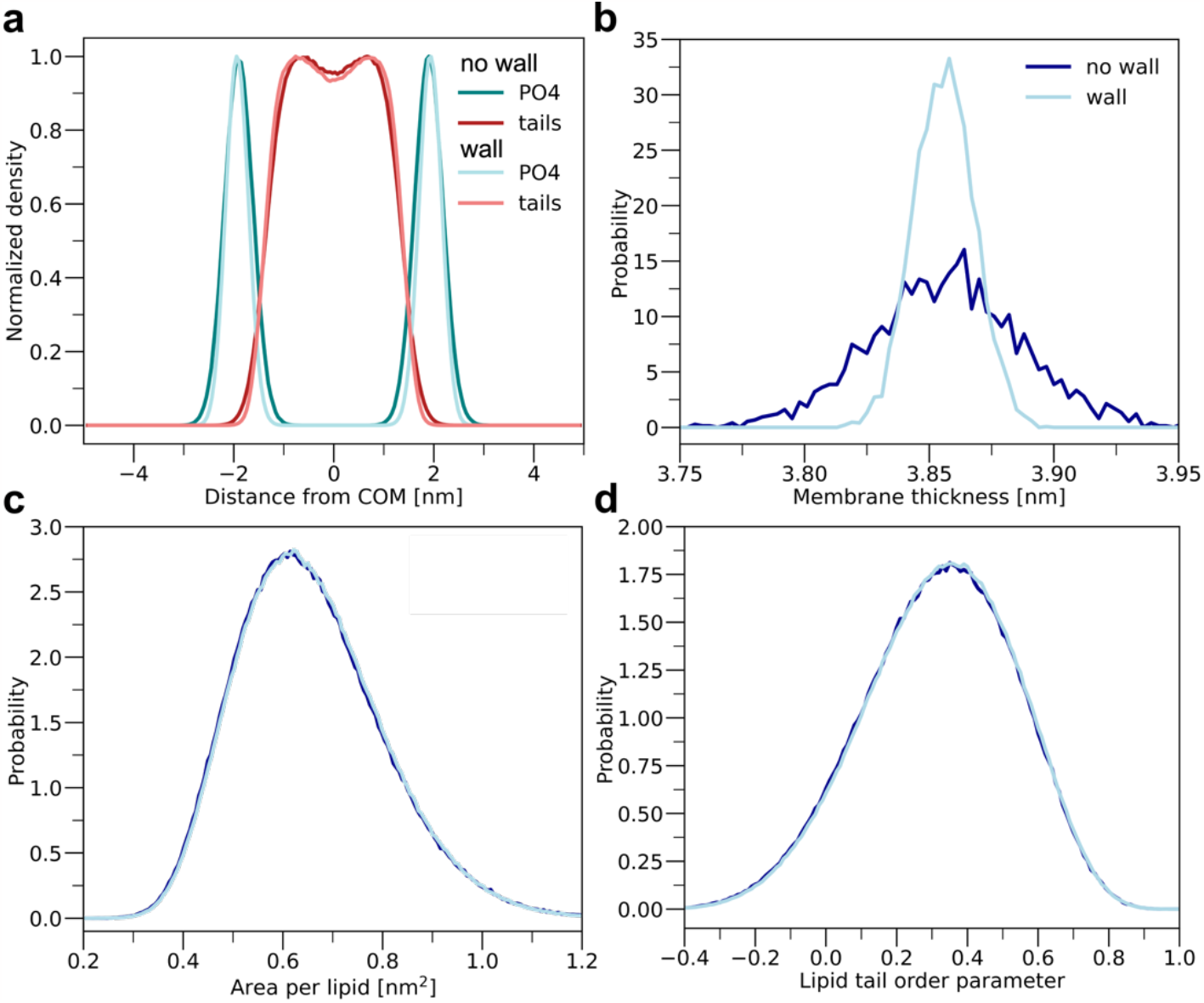
Comparison of membrane properties for a flat POPC bilayer with and without confining walls: (a) Density profiles of PO4 head groups (dark and light teal) and tail groups (red and coral) with respect to the bilayer center of mass (COM) and distributions of (b) membrane thickness, (c) area per lipid and (d) average lipid tail order parameter (dark blue and light blue). Darker colors denote simulations without wall beads, while lighter shades are used to indicate the presence of wall beads.

The influences of the wall beads on the lipid bilayer are negligible. For instance, the average positions of the lipid head and tails beads with respect to the bilayer center remain unaffected while their distributions become narrower as it is expected since the idea of the wall beads is to constrain them in the space (**Figure 1a**). The wall prohibits the protrusion of lipid head groups into the bulk water and thus all bilayer undulations, as can be seen from the difference in membrane thickness distribution in **Figure 1b**. However, removing all undulations but keeping the average membrane thickness is essential in order to apply a precise and uniform curvature gradient onto the membrane and analyze the curvature effects. The distributions of the area per lipid (APL) and average lipid tail order parameter are virtually indistinguishable (**Figure 1c and d**). Additionally, the lateral lipid diffusion of POPC in the presence and absence of wall beads is calculated as 6.2 ± 0.1 and 6.1 ± 0.2 x10^−7^ cm^2^/s, respectively. Taken together, we conclude that the wall has almost no influence on the structural and dynamical bilayer properties.

### 2.2 The sorting of cardiolipin depends on its protonation state, counterions, lipid environment, and concentration

Depending on their shape and degree of tail saturation, different lipids sense curvature to varying extents. In the following section, cardiolipin is used as a benchmark for our simulation setup due to its large negative intrinsic curvature of -1.1 nm^-1^ [19]. First, we only consider neutral CDL, hereinafter referred to as CDL0, carrying four mono-unsaturated tails (C20:1). The simulated membrane (**Figure 2a**) consists of two equivalent monolayers in terms of curvature and lipid composition, a 9:1 mixture of POPC/CDL0. The shape of the membrane mid-plane follows a sinusoidal function with an amplitude of 4 nm and a period of 20 nm (box length in x dimension), resulting in a broad range of mean curvatures from -1 to 0.1 nm^-1^. Simulating the system under constant area in the x-y plane and with periodic boundary conditions ensures the continuity of both leaflets, allowing the lipids to distribute according to their spontaneous curvature preference which is only limited by their in-plane diffusion.

**Figure 2.**
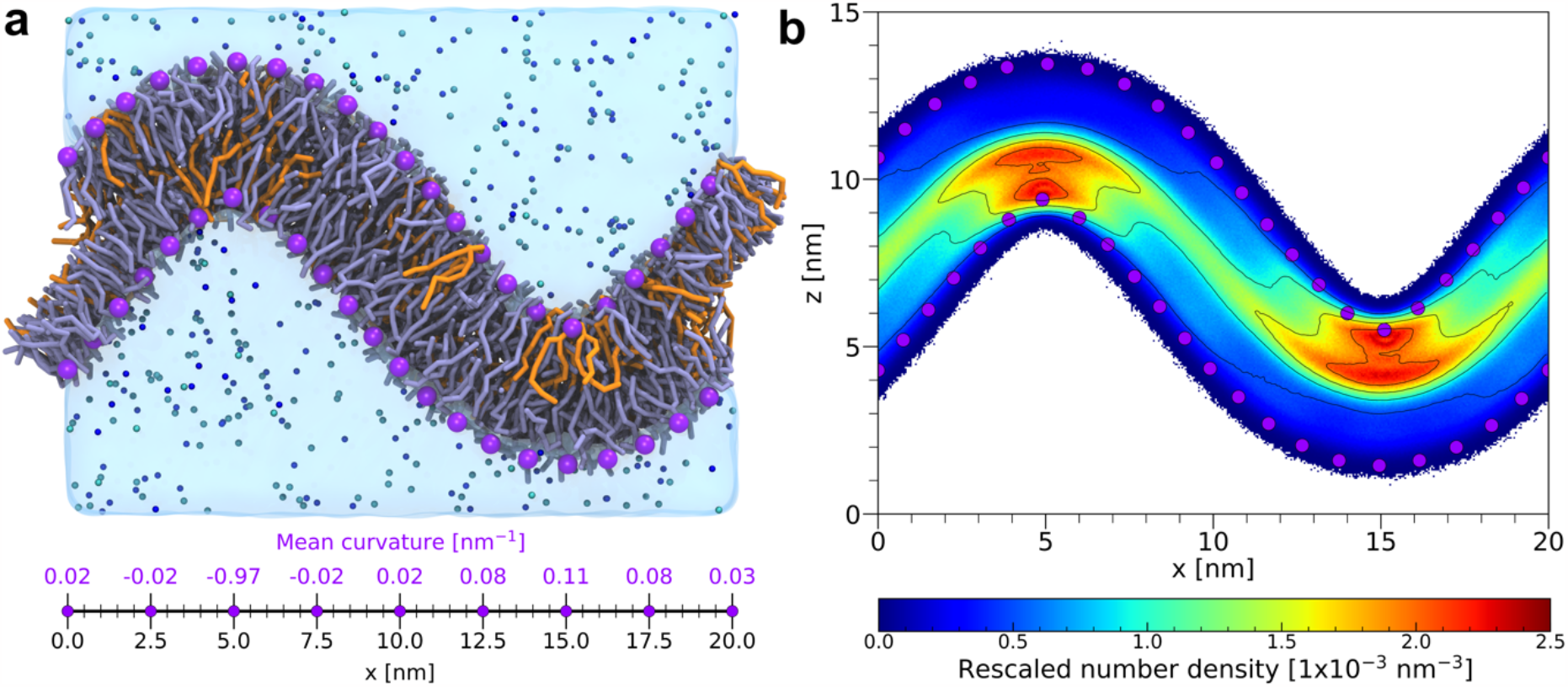
Distribution of POPC and CDL0 lipids in sine curved membrane. (a) Snapshot of the entire system (POPC/CDL0 9:1) after 5 µs of MD simulation showing CDL0 in orange, POPC in lilac, wall beads in purple, sodium and chloride ions in blue and cyan, respectively, and water in transparent blue. The axis shows the mean curvature of the lower leaflet along the x-dimension of the box. (b) The 2d number density of CDL0 lipids is computed as an average over the last 2 µs of 3 replicates. The density was measured using all CDL0 beads and normalized by the total number of CDL0 beads.

**Figure 2b** shows the 2d number density of all CDL0 beads averaged over the last 2 µs. The density of CDL0 is strongly increased at high negative curvature, centered at 5 nm and 15 nm for the lower and upper monolayer, respectively. The coneshaped nature of CDL0 greatly amplifies the lipid packing in regions with negative curvature which is in qualitative agreement with the lipid shape concept. It should be noted that the tail bead density of CDL0 in the midplane is overall increased as well due to the additional bead per tail compared to POPC.

Besides the molecular shape, the curvature preference of lipids can be influenced by their protonation state. CDL is a unique lipid with four acyl chains and two phosphatidyl headgroups, both can potentially carry a negative charge. However, the two phosphates groups are stereochemically nonequivalent [47] and their difference in p*K*_*a*_ has been discussed for many years due to possible implications on pathological processes such as Parkinson’s disease, Barth syndrome, and other cardiovascular diseases [48]. Recent studies [49-51] suggest that both phosphate groups are strong dibasic acids with similar p*K*_*a*_ values resulting in CDL being fully ionized at physiological conditions. Conversely, earlier FT-IR measurements [52-54] and DFT calculations [55] predict a large gap between the two p*K*_*a*_ values, indicating that the single anionic species predominates.

Here, both models are simulated, in the following denoted as CDL1 and CLD2, to assess the effect of the head group charge on the curvature-driven sorting behavior. For this, we measure the critical packing parameter (CPP) which is a quantitative measure of lipid shape, obtained from reference simulations of CDLs in flat POPC bilayers (see Supplementary Methods). While the size of the head group bead remains the same, the APL increases with an increasing head group charge, which leads to a reduction of the CPP (see Supplementary Table 1). A reduced CPP corresponds to a reduced intrinsic curvature. Based on the calculated CPP values, charged CDLs are considerably less cone-shaped than neutral CDL0, which finds its origin in the stronger hydration of the negatively charged CDL headgroups. The 2d lipid density in **Figure 3a-c** shows that all CDL models prefer negative curvature independent of the head group charge, however, the reduction in the CPP noticeably reduces the sorting behavior of both CDL1 and CDL2.

**Figure 3.**
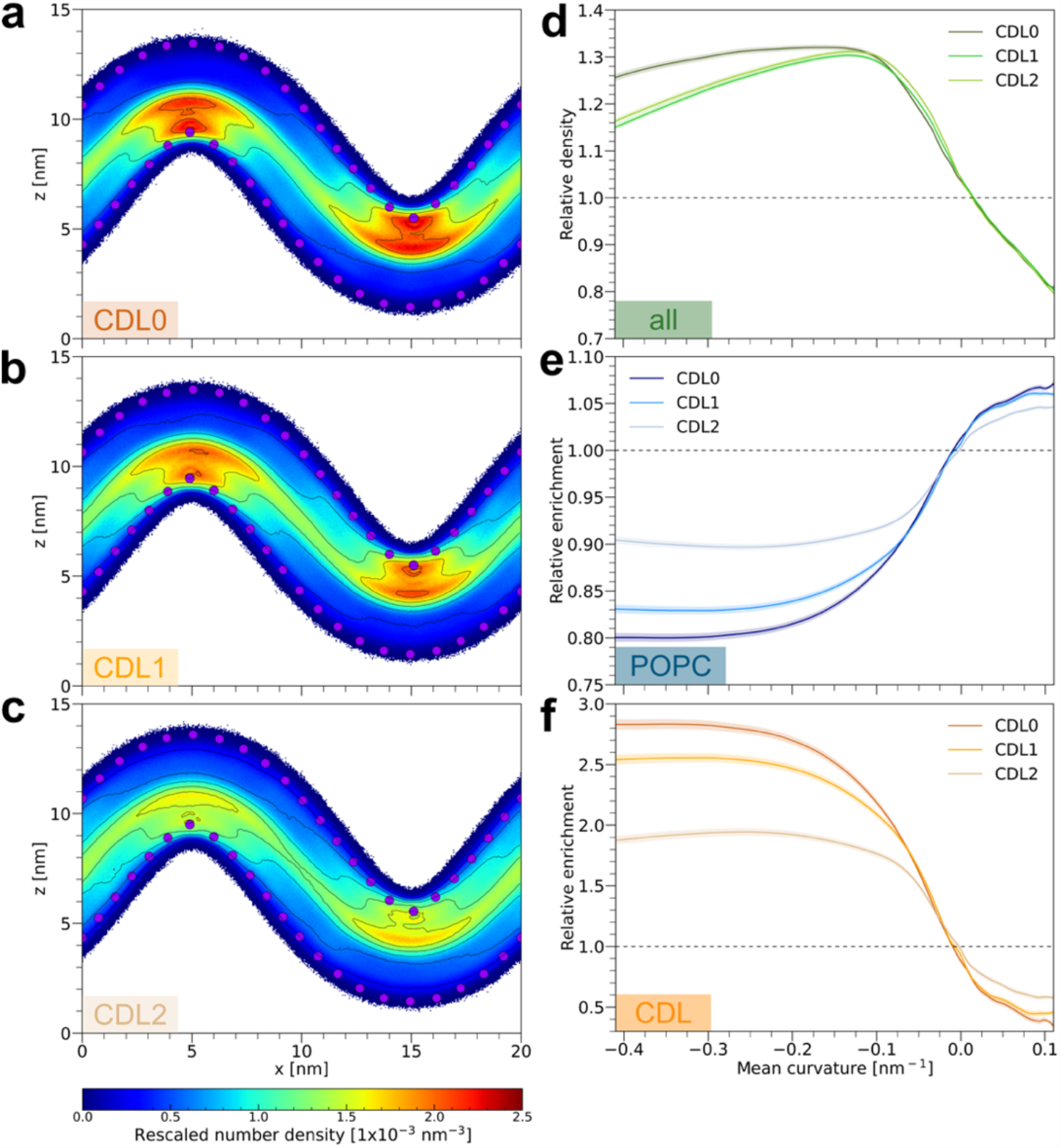
Increase in anionic charge of CDL reduces sorting preference. The 2d density of (a) CDL0, (b) CDL1, and (c) CDL2 is measured using all CDL beads and normalized by the total number of beads. (d) The relative density of all POPC and CDL lipids is calculated as the head group density of each species with respect to the frozen wall beads divided by the expected lipid density. The relative enrichment of (e) POPC and (f) CDL results from the relative lipid density of each lipid species dived by the combined relative density of all lipids (taken from d). The data of both leaflets are combined and fitted via Lowess regression. The shaded areas show 95% confidence intervals. Values above and below one (horizontal dashed line) indicate local enrichment and depletion, respectively.

We calculate the relative density (**Figure 3d**) and relative enrichment (**Figure 3e-f**) of lipid head groups versus the mean curvature to further elucidate lipid packing and distribution in curved membranes. The relative density, a measurement for the lipid packing, combines the lipid head group densities around the wall beads from both leaflets and is normalized by the expected lipid density assuming an equal distribution of lipids across each leaflet. Calculating the relative lipid density for a single lipid species and normalizing this density by the relative lipid density of all lipid species yields the relative enrichment (see Supplementary Methods). Taking all lipids together, the highest lipid density is observed in the region of negative mean curvatures with a maximum between -0.14 and -0.08 nm^-1^ (**Figure 3d**). Lipid head groups in this region are up to 32% more densely packed compared to flat regions. The opposite is true for positively curved regions, here the density decreases by up to 20%. The sorting behavior of the bulk lipid POPC is strongly influenced by CDL. Independent of the protonation state, CDL shows the strongest enrichment at high negative curvatures (< -0.20 nm^-1^), expelling POPC from these regions (**Figure 3e**). In positively curved regions, CDL is depleted to a varying extent depending on the head group charge. The larger the CPP, hence the more pronounced the cone-shaped character, the more enriched (depleted) the CDL lipids are in negative (positive) curvature (**Figure 3f**).

Unlike zwitterionic lipids like POPC, charged CDL lipids cannot shield themselves by aligning their head groups, which makes CDL very sensitive to the ionic composition of the surrounding aqueous phase. Interaction between anionic CDL and divalent cations, in particular, can significantly alter the physicochemical bilayer properties such as phase behavior [56], spontaneous curvature [57], and lipid packing [58], which has severe consequences on many cellular processes.

To probe the effect of Ca^2+^ on the sorting behavior of CDLs, we performed additional simulations with calcium instead of sodium chloride (at matching ionic strength). We find that the recent Martini 3 parameters correctly reproduce the strong clustering behavior of Ca^2+^ around anionic CDL, while no such clustering is observed for sodium ions (Supplementary Figures 1 and 2). The presence of divalent cations reduces the electrostatic head group repulsion, which subsequently induces the clustering of charged CDL lipids (see Supplementary Figures 2 and 3). **Figure 4** shows a substantial boost in the sorting behavior of CDL1 and especially CDL2 upon Ca^2+^ binding, even though the CPPs do not change significantly (see Supplementary Table 1). In the presence of Ca^2+^, CLD1 is up to 3.1 times enriched and CDL2 up to 3.4 times compared to flat bilayer regions. Interestingly, CDL0 (although it is neutral) also becomes more enriched. This result indicates that Ca^2+^ ions not only exert short-range attractive forces but also, in interplay with curvature, induce long-range clustering and sorting beyond what the CPP value would predict.

**Figure 4.**
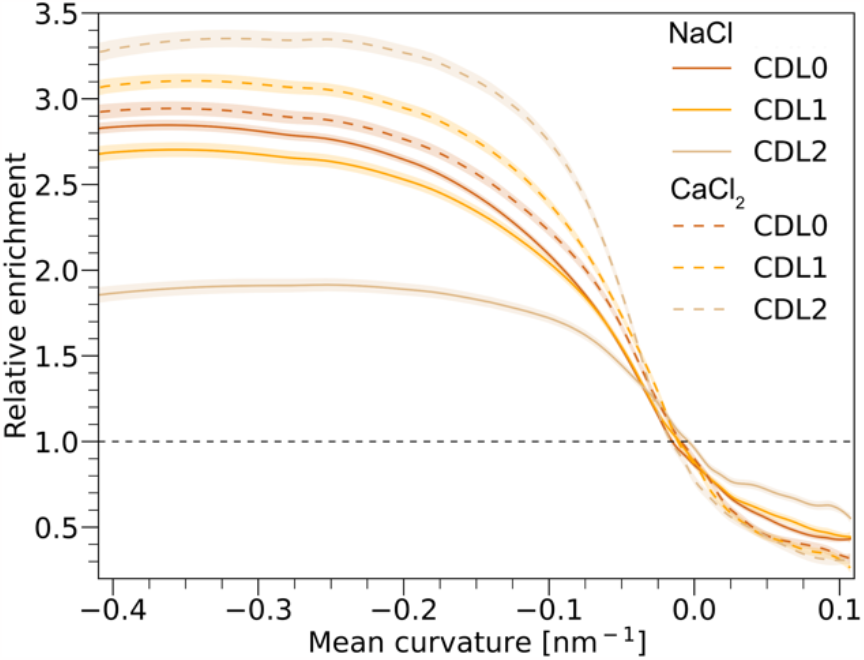
Ca^2+^ boosts the sorting behavior of CDL. Relative enrichment for the three different CDL protonation states in POPC in the presence of 150 mmol NaCl (solid lines) or 50 mmol CaCl_2_ (dashed lines). The data of both leaflets are combined and fitted via Lowess regression. The shaded areas show 95% confidence intervals. Values above and below one (horizontal dashed line) indicate local enrichment and depletion, respectively. All data is computed as an average over the last 2 µs for 3 replicas.

Next, we investigate the effect of the lipid environment on CDL shape and sorting considering different lipid head groups in the embedding membrane. In the following, we keep the 16:0-18:1 (PO) tails but focus on six different lipid head groups: zwitterionic PC, phosphatidylethanolamine (PE), and sphingomyelin (SM), anionic phosphatidylglycerol (PG), phosphatidylserine (PS) and di-anionic phosphatic acid (PA). In **Figure 5a** the bulk lipids are sorted according to the CPP values of the CDL lipids from flat reference simulations. In general, all CDL models appear most cone-shaped in a charged lipid environment except for POSM. The presence of the sphingosine chain makes the POSM tails asymmetric and slightly shorter than the regular fatty acid chains, which alters the lipid packing compared to other neutral lipids like POPC and POPE (see Supplementary Figure 6a). The effective shape of CDLs is least pronounced when surrounded by PE lipids, presumably because PE lipids themselves are coneshaped.

**Figure 5.**
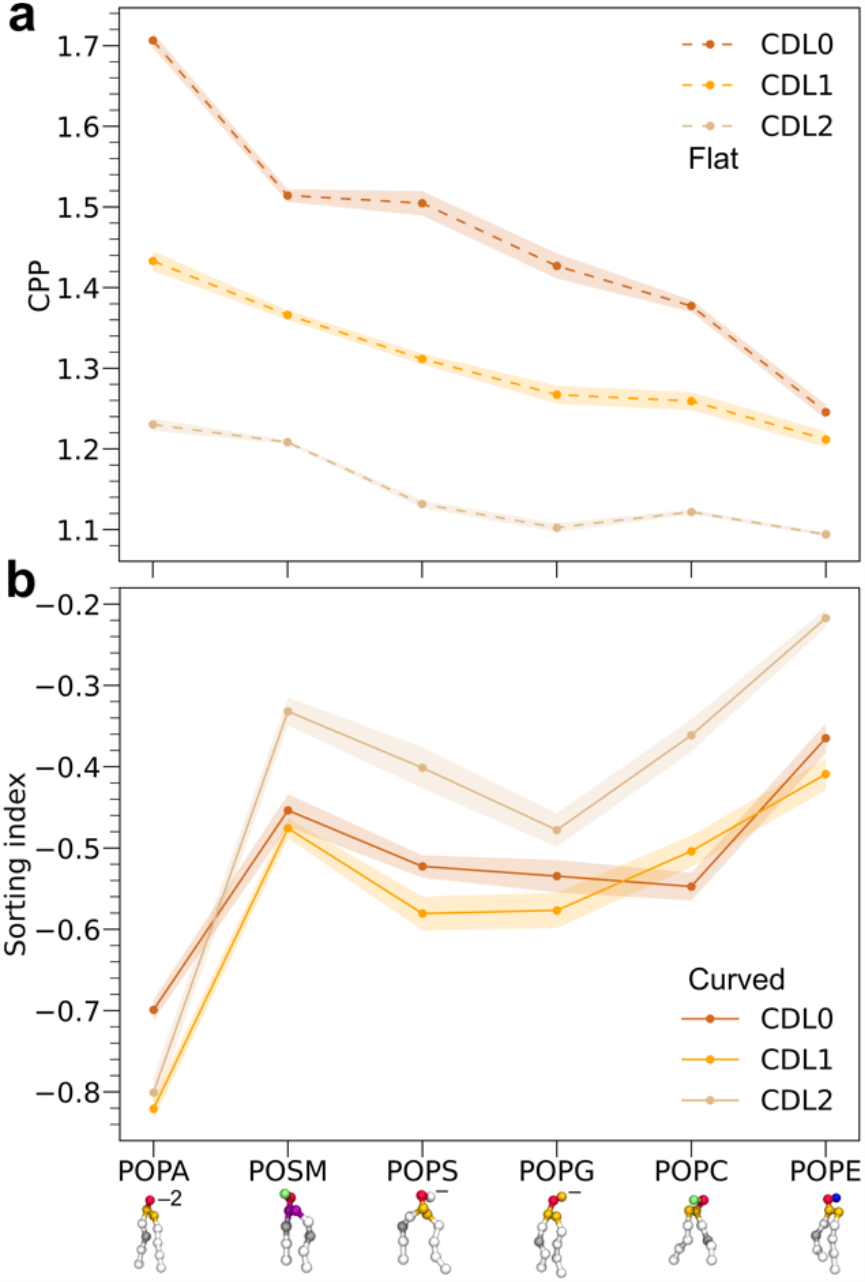
Surrounding lipids influence the shape and sorting of CDL. a) CPP and (b) sorting index for three different CDL protonation states in POPA (-2), POSM (±0), POPS (-1), POPG (-1), POPC (±0), and POPE (±0). CPP values for CDLs are computed from flat reference simulations using the same bilayer composition as curved systems, which are used to measure the sorting behavior. The sorting index quantifies the lipid density in positive vs. negative curvature regions normalized by the relative size of those regions. All data is computed as an average over the last 2 µs for 3 replicas. Standard errors are indicated as shaded areas.

To effectively compare the sorting behavior of the different CDL species, we introduce the sorting index, a measurement of lipid density in negatively vs. positively curved regions (see Supplementary Methods). A negative (positive) sorting index describes enrichment in negative (positive) curvature (**Figure 5b**). Overall, the sorting indices of the three CDL models increase with decreasing CPP values. Using di-anionic POPA as bulk lipids significantly enhances the localization of both charged and uncharged CDL lipids in negative curvature compared to all other lipids. Consequently, POPA lipids not only induce CDL clustering by reducing the electrostatic head group repulsion but also have a curvature preference on their own. POPA lipids show strong enrichment in positive curvature and therefore allow more CDLs to occupy negatively curved regions (see Supplementary Figure 6b). The opposite can be observed in POPE, which is known to prefer negative curvature [36, 43]. In this case, competition for negative curvature arises, which overall reduces the sorting of CDL independent of the protonation state.

Interestingly, CDL1 shows the strongest sorting behavior in most lipid environments despite CDL0 being far more cone-shaped than the charged CDL species. The limitations of the lipid shape concept become even more clear for CDL lipids in POSM. Despite the large CPP values, the sorting behavior of all three CDL models is greatly reduced in POSM. This result indicates that the effective lipid shape is not a static property obtained from flat reference simulations. It rather depends on available curvature and optimal packing of all the lipids that are present in the curved regions.

Not only the protonation state, counterions, and lipid environment can affect the properties of CDL-containing bilayers but also their concentration has previously been shown to greatly influence membrane structure and dynamics [37]. The CDL concentration in the IMM ranges from 5 to 20%, however, CDL is also present in the outer mitochondrial membrane (OMM) but at a much lower concentration (1-5%) [59]. CDL plays an important role in the regulation of the OMM permeabilization, a crucial event in the intrinsic apoptosis pathway. By recruiting tBID to the contact sites between OMM and IMM [60-62], the CDL concentration locally increases up to 25% [63]. Furthermore, the overall concentration of CDL in the OMM increases during apoptosis leading to the formation of CDL-enriched microdomains, which influences the mitochondrial membrane fission [64].

Given this broad range of CDL concentrations in biologically relevant systems, we investigate the concentration-dependent shape and sorting behavior of CDL (**Figure 6**). For all CDL protonation states, the CPP decreases with increasing concentration due to packing constraints arising from their geometrical shape (**Figure 6a**). Even at high CDL concentrations, neutral CDL0 remains slightly more cone-shaped than anionic CDL1 and CDL2. Following their CPP values, all CDL models display a high negative sorting index which increases with increasing concentration, though still maintaining a clear preference for the negatively curved membrane regions. As CDL prefers the hexagonal over the lamellar phase, packing the four bulky lipid tails into the curved bilayer becomes increasingly problematic at higher concentrations therefore sufficient POPC is needed to stabilize the lamellar phase. Even in the absence of electrostatic repulsion between the head groups, a small increase in concentration drastically reduces the observed sorting behavior. At low CDL concentrations (< 20%) the sorting index strictly follows the order of CPP values (CDL0 > CDL1 > CDL2). At higher concentrations, however, the way the lipids are packed in negative curvature regions changes (see Supplementary Figure 4), which allows for better packing of CDL1 at very high concentrations despite the electrostatic head group repulsion. Interestingly, no such effect could be observed for CDL2, suggesting that an asymmetric charge distribution might be most beneficial for the packing of CDLs at high concentrations.

**Figure 6.**
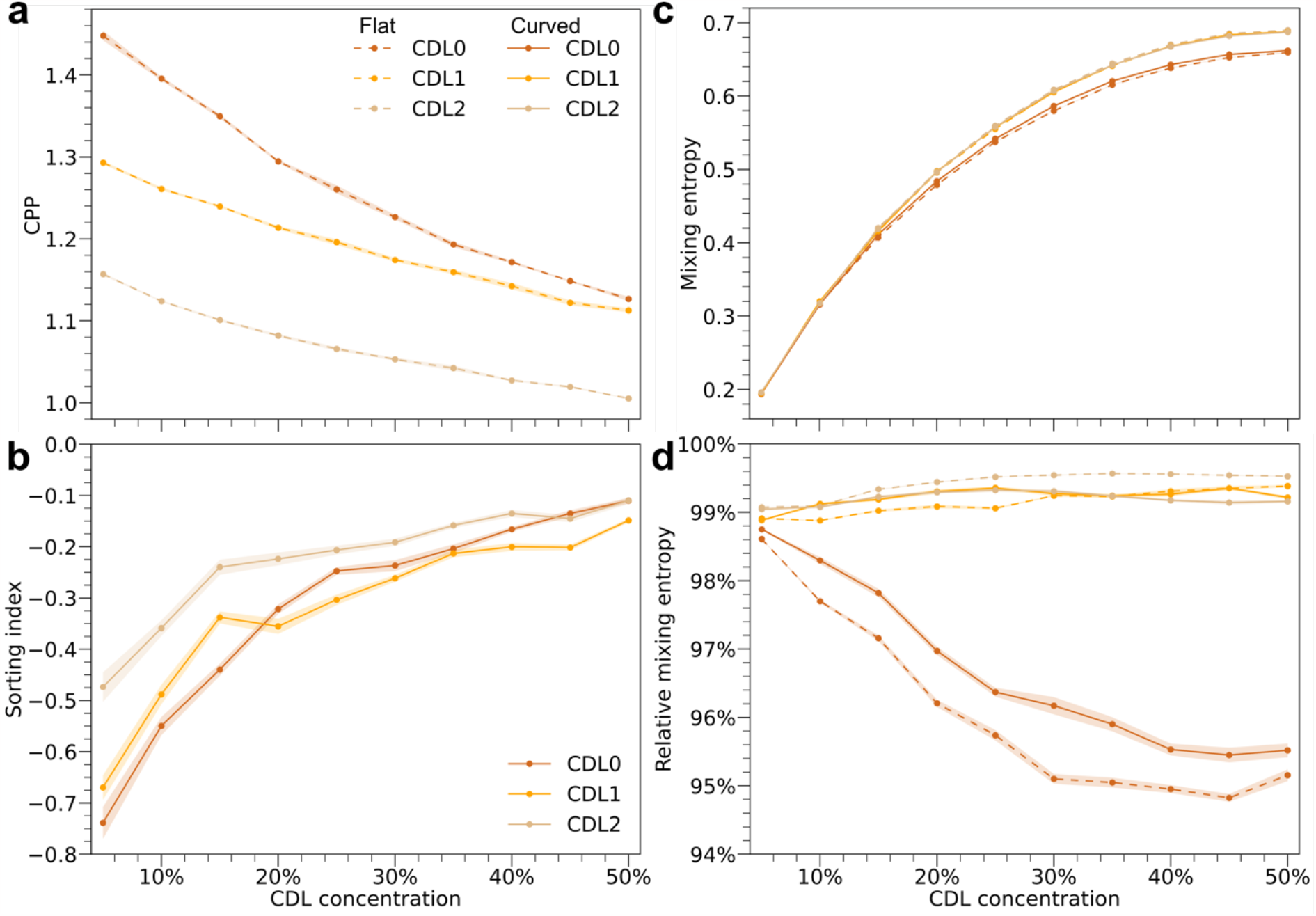
Increase in CDL concentration reduces cone-shaped character, sorting behavior, and mixing entropy. (a) CPP, (b) sorting index, (c) absolute, and (d) relative conditional mixing entropy for the three different CDL protonation states at ten different concentrations (5-50%) in POPC. The sorting index quantifies the lipid density in positive vs. negative curvature regions normalized by the relative size of those regions. All data referred to as ‘Flat’ (dashed lines) is computed from flat bilayer simulations with the same lipid ratios as ‘Curved’ systems (solid lines). All data is computed as an average over the last 2 µs for 3 replicas. Standard errors are indicated as shaded areas. (Relative)

### 2.3 Cardiolipin sorting does not work against mixing entropy

Besides the effect of the altered lipid packing, an increase in CDL concentration also causes a rise in lipid mixing entropy, which is generally assumed to counteract lateral lipid segregation and should therefore reduce the sorting behavior. A theoretical model by Derganc [23] predicts that the energetic benefit from curvature matching, which relaxes membrane curvature stress, is mostly balanced by the homogenizing effect of the mixing entropy. The author concludes that curvature-driven lipid segregation in Golgi cisterna is effectively limited and can only be enhanced when lipids aggregate into larger clusters with non-zero spontaneous curvature.

To estimate the effect of lipid mixing on curvature-induced sorting of CDL, we calculate the conditional mixing entropy [65] using a nearest neighbor approximation (see Supplementary Methods). The conditional mixing entropy describes the probability of finding an individual lipid in all possible lipid environments. Consequently, the ratio of lipids in the membrane defines the theoretical maximum mixing entropy. **Figure 6c** shows that the mixing entropy increases with increasing CDL concentration for both flat and curved membranes, correlating with the decrease in the sorting index. This indicates that the mixing entropy limits the sorting of CDL lipids at high concentrations. However, it appears the presence of curvature and thus curvature-induced sorting of CDLs does not reduce the mixing entropy compared to the flat bilayer (**Figure 6c**). For a better comparison, the mixing entropy for each concentration is normalized by the respective theoretical maximum mixing entropy (**Figure 6d**). Surprisingly, we find that the relative mixing entropy for curved bilayers is slightly increased for a wide range of concentrations for CDL0 and CDL1. The relative mixing entropy demonstrates that CDL and POPC lipids in curved membranes are often more ideally mixed than in the flat reference system. This effect is most pronounced for CDL0, showing an increase of 1% in relative mixing entropy, but still visible for a wide range of concentrations for CDL1 and CDL2.

Lipid sorting increasing the mixing entropy may seem contradictive because we see enrichment of CDL in the negative curvature regions (see **Figure 3f**). This implies that overall, the system gains more order. But contrary to current beliefs, the packing of lipids with non-cylindrical shapes is not ideal in flat membranes either, thus the sorting of lipids in curvature gradients could reduce the packing stress. As seen in **Figure 3d** the lipid density is not uniform but increased in negative curvature regions. This does not only apply to CDL but also to POPC lipids, which enhances the lipid packing in those regions. This means that not only more CDL lipids are surrounding each other but also more POPC lipids are in the neighborhood of CDLs. To confirm this hypothesis, we computed the RDFs for both lipid components in flat and curved systems (see Supplementary Figure 5). These RDFs clearly show an increased first and second neighbor peak. We hypothesize that this effect of increased local packing leads to an overall similar mixing behavior compared to the flat reference systems. Given the conditional entropy as a metric for lipid mixing, these results show that arguments on the mixing of lipids in curved systems are more complex than generally assumed. The global reduction in entropy due to curvature-driven sorting could be less severe considering the increased local mixing.

### 2.4 Cooperativity is not required for cardiolipin sorting

Generally, lipid sorting is assumed to be a cooperative effect requiring lipid-lipid or lipid-protein interactions to offset the costs of the mixing entropy. Huang *et al*. [66] estimated an energy difference of 0.01 *k*_*B*_*T* for the curvature matching of a single lipid with an intrinsic curvature of 0.1 nm^-1^ in a rodshaped cell with a polar hemisphere curvature of 0.004 nm^-1^. The authors conclude that in the presence of thermal fluctuations, the curvature preference is insufficient to cause lipid segregation.

This finding is in line with other studies [13, 23, 25, 28, 31], however, most of this work does not focus on lipids with high intrinsic curvature like CDL and only considers relatively small curvatures (radii of curvature >20 nm).

To examine if a single CDL molecule can effectively sort in a highly curved bilayer, mimicking the curvature gradient which can be found in the inner mitochondrial membrane (IMM) (-2 < *H* < 10 nm^-1^), we use umbrella sampling along the curved membrane to compute the PMF of curvature sorting. **Figure 7a** shows the PMF as a function of the x-position with a free energy minimum centered at 5 nm for all three CDL protonation states, the region of maximum negative curvature. Although the energy difference between the opposite curvature regions is small, in the order of 3-4 *k*_*B*_*T* (**Figure 7b**), the curvature preference of a single CDL molecule based on its molecular shape is evident. In agreement with their CPP values, CDL2 shows the least preference for negative curvature compared to the more cone-shaped CDL1 and CDL0.

**Figure 7.**
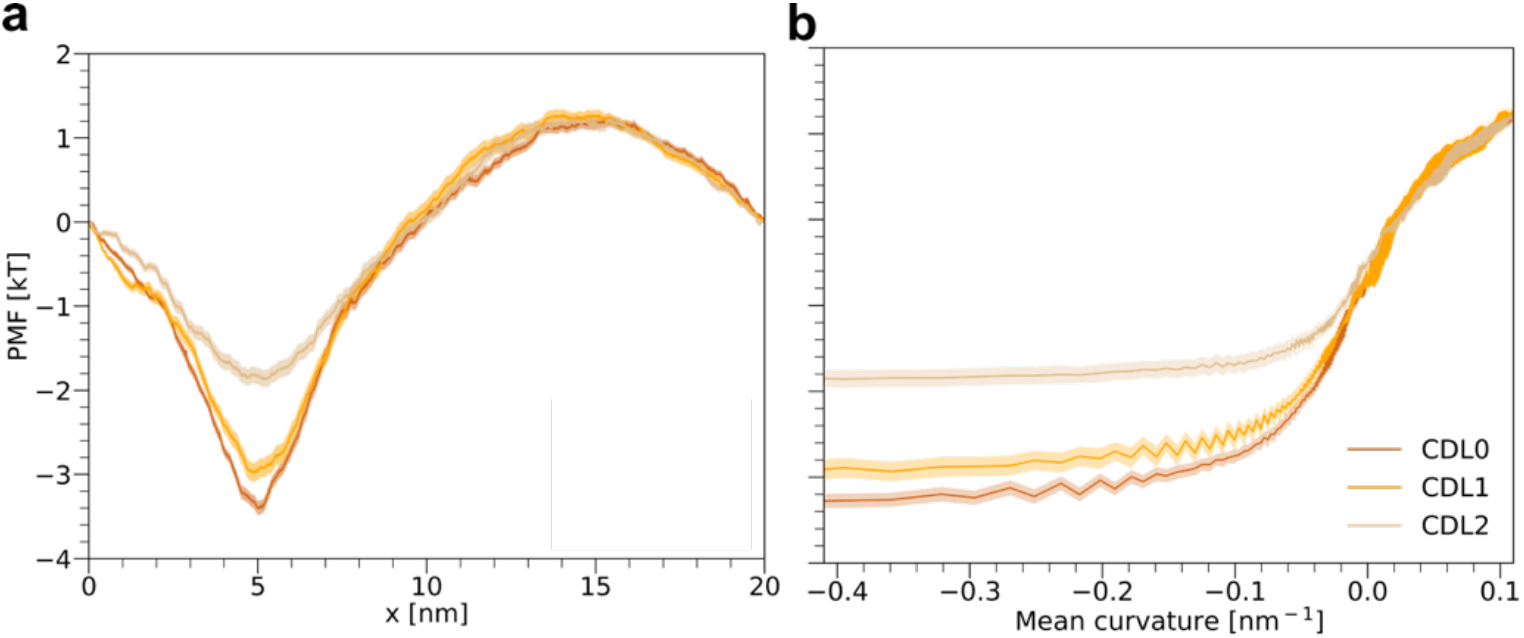
PMF of curvature-mediated CDL sorting. PMF of moving a single CDL molecule across the sine curved bilayer for all three different CDL protonation states is shown as a function of x-position (a) and corresponding mean curvature (b). The shaded areas correspond to the standard error computed by bootstrapping.

### 2.5 Lipid sorting depends on tail length, degree of saturation, and head group type

So far, we demonstrated that curvature can induce substantial lateral segregation in cardiolipin-containing bilayers due to their prominent cone-shaped character. Mostly the CPP value correlated well with the observed sorting behavior apart from certain limitations e.g., charged CDL species and presence of divalent ions.

In the following section, we will systematically investigate the curvature preferences of various other phospholipids. To simplify the interpretation, we will again focus on simple binary mixtures in which 10 mol% of the target lipid is embedded in a bulk lipid membrane. We will disentangle the effect of (i) tail length by simulating PC lipids with 3 to 6 tail beads which correspond to 8 to 26 carbon per tail; (ii) tail saturation using PC lipids with different degrees of saturation from fully saturated to polyunsaturated; and (iii) head group using PA, SM, PS, PG and PE lipid types. To exclude the effect of chain interdigitation when increasing tail length and saturation, the bulk lipid used in these simulations is DNPC (di-C26:1). Simulations concerning the effect of variations in the head group use POPC as the bulk lipid.

The results of these simulations are shown in **Figure 8**. Again, we compare the lipid shape expressed by the CPP value with the observed curvature preference. While short PC lipids like DTPC and DLPC are considerably cone-shaped, increasing the tail length drastically decreases the CPP. Despite the similarities in shape between PC and CDL lipids, PC lipids experience the opposite sorting trend. The more pronounced the cone-shaped character, the more depleted (enriched) the PC lipids are in negative (positive) curvature (**Figure 8a**). Packing of long cylindrical DNPC lipids into a curved bilayer creates significant gaps in positive curvature regions, thereby exposing the lipids’ tails to the hydrophilic environment. Filling these gaps with shorter lipids optimizes the packing and is in this case more beneficial than the segregation of those lipids to their intrinsically preferred curvature region. The longer the lipid tails, the less efficiently those gaps can be filled and the apparent sorting to positive curvature decreases.

**Figure 8.**
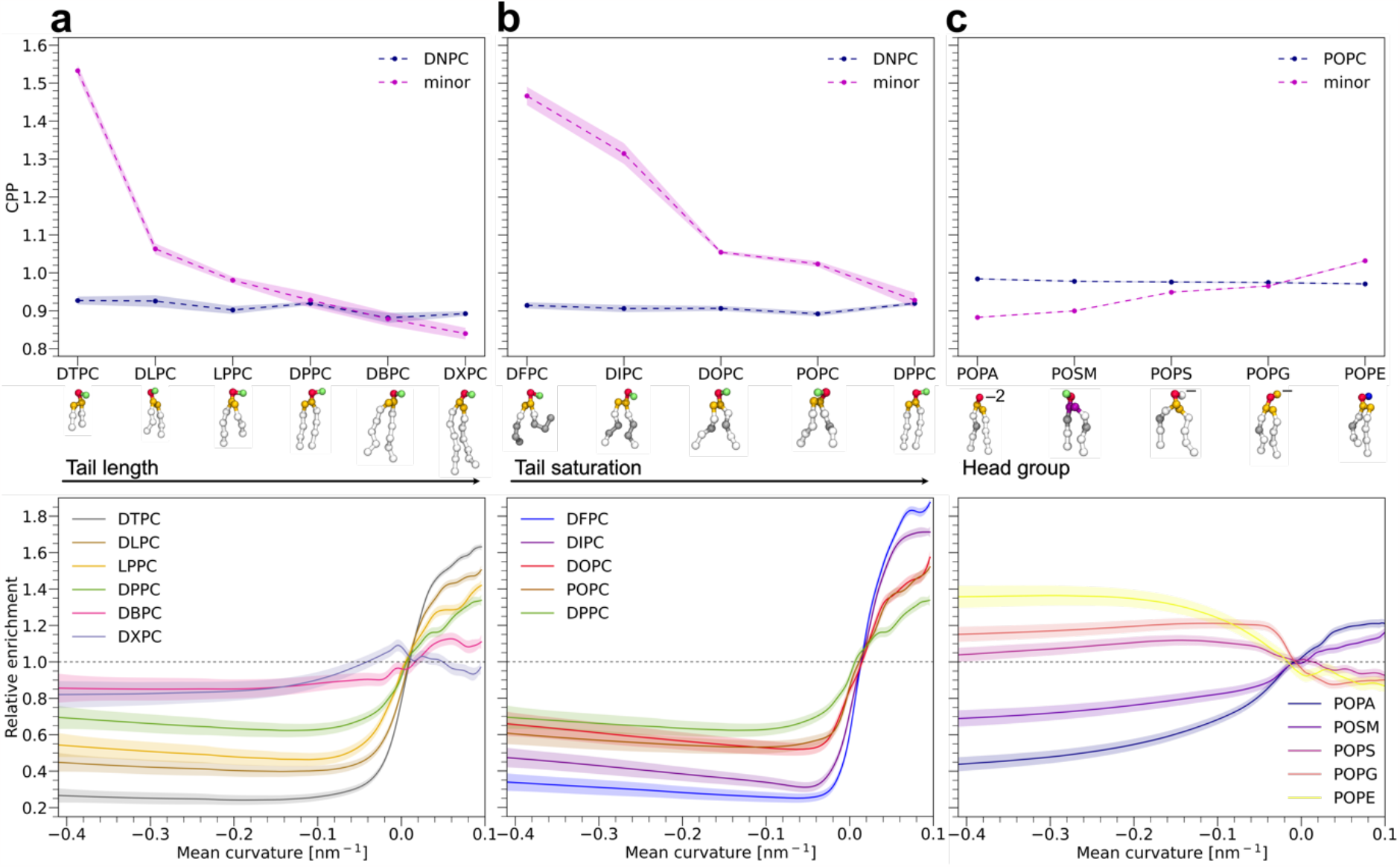
Effect of (a) tail length, (b) degree of saturation, and (c) lipid head group type on shape and sorting behavior. CPP (top) and relative enrichment (bottom) of various lipids in (a-b) DNPC (di-C24:1) or (c) POPC in the presence of 150 mmol NaCl. CPP values are computed from flat bilayer simulations with the same lipid ratios as the curved system used to analyze the sorting behavior. All data is computed as an average over the last 2 µs for 3 replicas. Shaded areas indicate standard errors (top) or 95% confidence intervals (bottom).

The degree of tail saturation influences the sorting behavior in a similar manner. Highly unsaturated lipids like DFPC (di-C16:3) and DIPC (di-16:2), which have a negative intrinsic curvature preference according to their CPP values, show significant sorting to positive curvature. While all lipids in **Figure 8b** have the same number of tail beads, the degree of saturation changes the effective tail length, making (poly)unsaturated lipids significantly shorter than saturated ones. Packing of (poly)unsaturated lipid in positive curvature is, therefore, most effective the lower the degree of saturation is. Furthermore, the magnitude of the packing-induced sorting does not correlate with the CPP values. Despite, DTPC and DFPC having very similar shapes in flat bilayer simulations, polyunsaturated DFPC is up to 25% more enriched in positive curvature regions than fully saturated DTPC. Polyunsaturated lipids like DFPC have highly flexible tails which can perfectly adjust their shape to adapt to their surroundings, which cannot be captured by their intrinsic shape.

Conversely, for different types of head groups, we find good agreement between lipid shape and curvature-driven sorting (**Figure 8c**). Cone-shaped POPE lipids show noticeable segregation to negative curvature, consistent with experimental reports [67, 68] and the above observed curvature competition between CDL and POPE. Likewise, inverted cone-shaped lipids like POPA and POSM, which have a lower CPP value compared to the bulk lipid POPC, experience strong to moderate depletion from negatively curved regions. Despite the small difference in CPP, di-anionic POPA is more enriched in positive curvature than POSM. Monoanionic POPS and POPG display moderate enrichment at negative curvatures, even though they have CPP values close to one, which indicates a cylindrical shape. As shown above for charged CDL models, electrostatic head group repulsion can alter the intrinsic curvature preference. To test the influence of the protonation state, we reduced the anionic charge POPA, which drastically changes the curvature-induced sorting, leading to enrichment in negative curvature similar to POPS and POPG (see Supplementary Figure 8).

In the presence of divalent ions, however, the sorting behavior of anionic lipids can be modified as we already demonstrated for CDL above. Recent studies [69-73] have shown that Ca^2+^ ions trigger aggregation of negatively charged lipids like PS and phosphatidylinositol-4,5-bisphosphate (PIP2) which induces spontaneous curvature. However, the direction of induced curvature is currently still under debate. In the present work, the addition of Ca^2+^ causes minor enrichment of mono-anionic PS and PG lipids in regions dominated by negative curvature, while di-anionic POPA favors neutral curvature in the presence of Ca^2+^ (see Supplementary Figure 7). Lipid clustering in both flat and curved bilayers is induced by the formation of Ca^2+^ bridges between negatively charged phosphate groups (see Supplementary Figure 9). However, more Ca^2+^ ions are sequestered by anionic lipids in curved bilayers, especially by di-anionic POPA, which leads to the formation of significantly larger lipid clusters. Consequently, larger lipid clusters show stronger curvature preference.

Although the presented results focused on symmetric lipids mixtures, we found reasonable agreement with earlier works [39, 43, 44], studying the curvature-driven sorting of lipids in complex asymmetric mixtures. Minor differences, for example in the sorting of polyunsaturated lipids, can be explained by competing curvature preferences of head and tail groups. It should be noted, that also non-intuitive sorting might arise from differential stress [74] in those asymmetric mixtures, which could overrule the intrinsic curvature preference of individual lipids.

### 2.6 Interleaflet coupling, phase separation, and line tension can offset sorting preferences

Biological membranes, in particular plasma membranes, are heterogeneous entities. However, there is still no consensus on the mechanisms that control lateral organizations of cellular membranes. Several mechanisms have been proposed such as phase separation due to the miscibility of its components (lipid rafts)[75], specific lipid-lipid interactions [76], membrane curvature [77], and interleaflet coupling [78].

Next, we consider the behavior of different binary lipid mixtures containing highly polyunsaturated DFPC lipids. In the presence of other (poly)unsaturated bulk lipids like DIPC and DOPC, no phase separation occurs and DFPC lipids experience minor enrichment in positive curvature due to the improved packing of the polyunsaturated tails (**Figure 9a**). In the presence of fully saturated DPPC and single unsaturated POPC, however, DFPC lipids become enriched in both positive and negative curvature and almost entirely depleted in zero curvature (see Supplementary Figure 10a). This simultaneous enrichment cannot be explained by the molecular shape. Here, non-intuitive sorting behavior arises from the strong interleaflet coupling of the DFPC-rich phase and the promiscuous character of highly flexible and adaptable polyunsaturated tails.

**Figure 9.**
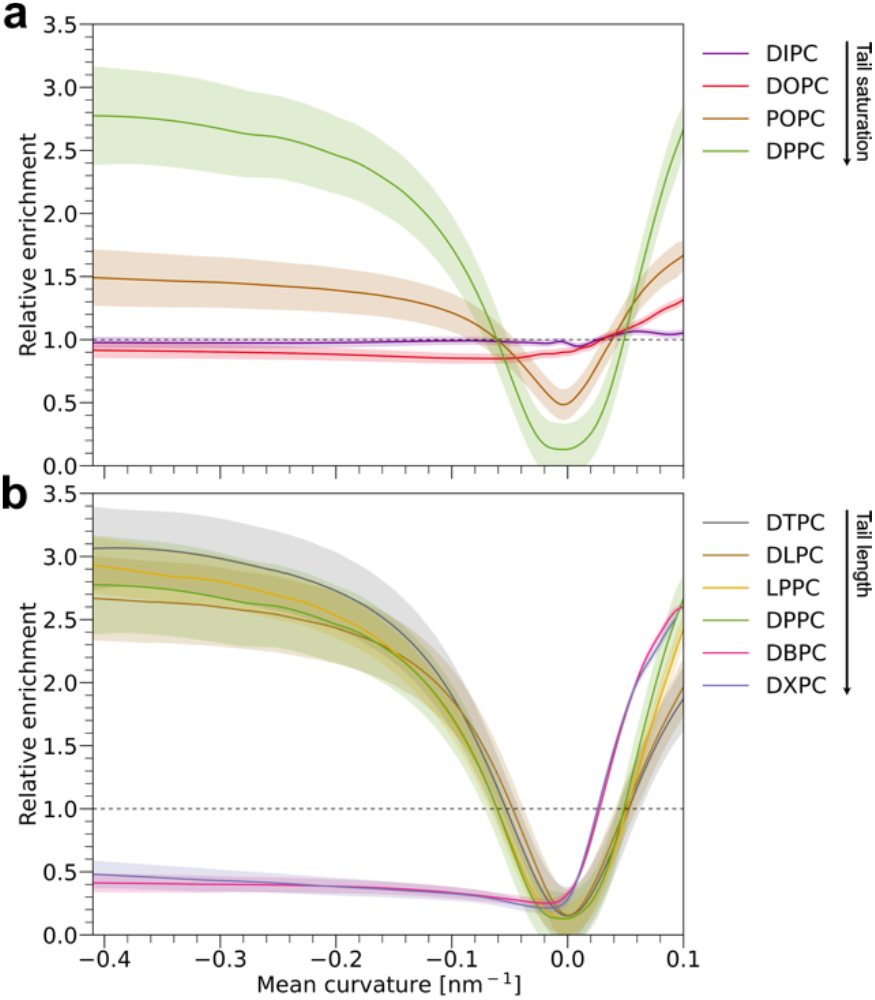
Nanodomain formation in binary mixtures of DFPC and PC lipids influences DFPC sorting. Relative enrichment of DFPC in PC lipids with (a) varying degrees of tail saturation and (b) tail length. Shaded areas indicate 95% confidence intervals.

To further examine the power of interleaflet coupling, the sorting preference of DFPC in mixtures with fully saturated PC lipids and varying tail lengths is measured in **Figure 9b**. In all mixtures, DFPC shows stronger enrichment compared to simulations of DFPC in single unsaturated DNPC (**Figure 8b**). Remarkably, only in the presence of short lipids, i.e., with a lipid tail length of 4 beads or less (C8:0-C16:0), DFPC lipids are enriched in both positive and negative curvature. This indicates that the tail beads of the formed nanodomains need to be in proximity to the bilayer midplane to effectively couple the lipid domains in both leaflets.

Last, we demonstrate how lipid curvature preference can overrule interleaflet coupling, phase registration, and line tension contributions. We considered two different ternary mixtures, one containing DPPC, CHOL, and DFPC and the other containing DPPC, CHOL, and DIPC, each in a molar ratio of 0.35/0.35/0.3. The DIPC system is less strongly phase-separated due to the reduced level of tail unsaturation compared to DFPC, as can be seen from the difference in lipid mixing entropies in Supplementary Figure 11. Nonetheless, both systems show visible phase separation into CHOL-enriched *L*_*o*_ and CHOL-depleted *L*_*d*_ phases (**Figure 10a, b**). In agreement with experimental studies on GUVs [27, 28] we find, that the highly phase-separated DPPC/CHOL/DFPC mixture shows alignment of the thicker and stiffer *L*_*o*_ phase in flat bilayer regions (**Figure 10a, c**). Due to the chosen molar ratios and system size, the *L*_*o*_ phase is sandwiched between the maximum curvature regions. If curvature sorting would be dominating, we expect two non-continuous *L*_*d*_ phases in each leaflet segregating to bilayer regions of positive and negative curvature, respectively, resulting in four separate lipid phases. While this scenario would maximize the curvature matching of lipid phases, the energetic contribution of the line tension at the interface of *L*_*o*_ and *L*_*d*_ phases cannot be neglected. Doubling the interfacial area between coexisting phases would also double the line tension, which has been proposed to drive membrane deformation, fission, and budding [79]. Although curvature preference might not be the most dominant factor for the lateral distribution of lipid phases, Baumgart and coworkers [80] have shown how line tension and curvature coupling can work in concert to generate long-range domain ordering. They revealed that factors such as temperature and membrane composition can strongly alter the appearance of coexisting phases. Similarly, we find that increasing the concentration of DPPC and CHOL leads to the expansion of the *L*_*o*_ phase, consequently pushing the *L*_*o*_ phase further into highly curved regions. To reduce the energy penalty associated with the bending of the *L*_*o*_ phase in regions of high curvature, two metastable *L*_*d*_ phases appear which align at maximum curvatures in both leaflets (Supplementary Figure 12). Due to this particular membrane composition, the line tension is reduced and curvature preference can partially overcome the phase-separating effect of the line tension. As a consequence of the strong phase separation in the DPPC/CHOL/DFPC system, the stiffer CHOL-rich *L*_*o*_ domains show elevated densities around zero curvature and are highly correlated in both leaflets (**Figure 10a, c**). Domain registration in phase-separated systems has been extensively studied (see review [81]) and can be modulated by lipid tail interdigitation, cholesterol flip-flop, membrane undulations, and line tension. A recent theoretical study [82] predicts that line tension alone is sufficient to drive domain registration, although the efficiency decreases with increasing domain size.

**Figure 10.**
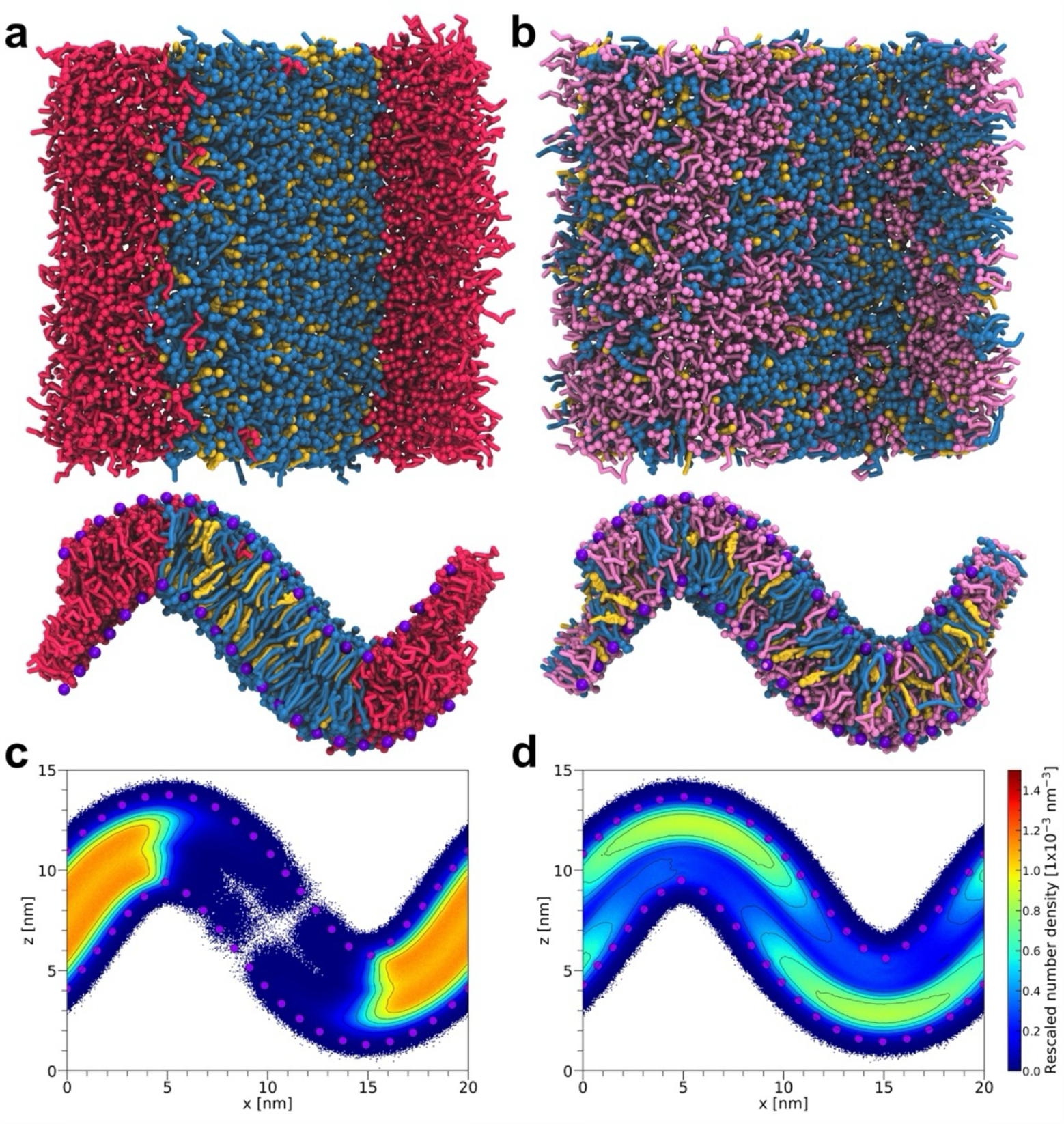
Curvature influences lateral segregation of phase-separated ternary lipid mixtures. Top and side view of (a) DPPC/CHOL/DFPC (0.35/0.35/0.30) and (b) DPPC/CHOL/DIPC (0.35/0.35/0.30) mixtures in curved bilayers after 10 µs of simulation. Lipids are colored as follows: DPPC - blue, DFPC - red, CHOL – yellow, and DIPC - pink. Wall beads in top-view images are omitted for clarity. The 2d number density of (c) DFPC and (d) DIPC lipids, corresponding to the *L*_*d*_ phase in each system, is computed as an average over the last 2 µs of 3 replicates.

It was found that the energetically most favorable configuration is anti-registered domains, where the line tension completely vanishes. While domain anti-registration has so far only been observed *in silico* [82-84] because it requires equally sized *L*_*d*_ and *L*_*o*_ domains, the effects of line tension and domain registration might play an important role in signal transduction across the plasma membrane of cells [85]. Decreasing the hydrophobic thickness mismatch between the *L*_*o*_ and *L*_*d*_ phase by using DIPC (di-C16:2) instead of DFPC (di-16:3), could in principle promote domain anti-registration. However, we only find domains being registered across the two leaflets in flat reference simulations (Supplementary Figure 13). Surprisingly, applying curvature to the weakly phase-separated DPPC/CHOL/DIPC system leads to the formation of anti-registered domains with the DIPC-rich *L*_*d*_ phases being located in regions with positive curvature (**Figure 10b, d**). Even when starting from biased configurations with all DIPC lipids in perfect registration or inverse antiregistration (all DIPC lipids placed in negatively curved regions), the positive curvature anti-registration of the *L*_*d*_ phase can be restored. Consequently, the CHOL-rich *L*_*o*_ domain favors negative curvature, consistent with simulations of CHOL in a binary mixture with POPC (Supplementary Figure 13) and the previously reported negative intrinsic curvature of CHOL [86, 87]. This result demonstrates the remarkable ability of membrane curvature to alter the phase behavior of complex membranes.

## 3. Discussion and conclusion

### 3.1 Pros and cons of the scaffolding approach

Despite curvature being an essential property of all biological membranes, simulating realistic membrane shapes has only recently become computationally feasible [33]. Although it is already possible to simulate the membranes of an entire organelle [88], available time scales are still very limited and therefore require a multiscale approach to provide a realistic picture of biomembranes. To efficiently study the influence of curvature on the underlying membrane structure and dynamics, various methods are available including membrane buckling by lateral stress [35, 36, 39, 89], applying a harmonic potential [90, 91], the use of solid support [41, 92] or scaffolding particles [93, 94]. As discussed in a recent review [95], the latter approach, which has been used here, comes with a certain number of limitations. Most importantly, flat reference simulations are needed to determine the bilayer thickness before setting up curved systems. This might lead to problems when simulating mixed bilayers with significant differences in tail length or strong phase separation. Additionally, the change in lipid density in curved regions alters the overall lipid packing, which requires careful control of the number of lipids when using a fixed membrane area. To realize a tensionless membrane and avoid artifacts at the periodic boundaries an iterative procedure has been used to match the membrane area and the number of lipids (see Supplementary Methods). Despite these limitations, the wall bead method is a simple way to set up curved bilayers, which comes with significant advantages over other methods. First, it allows precise control of arbitrary curvature gradients, while harmonic potentials can only apply a uniform curvature. Other methods like membrane buckling can only indirectly control and approximately sustain the imposed curvature. Second, wall beads can also be used for straightforward analysis of curvature-induced sorting. Although the bilayer thickness must be known a priori, it can easily be adjusted to accommodate various lipid phases with different heights by matching the maximum height. Furthermore, curved bilayers could potentially be combined with flat, unrestrained reservoirs to allow the bilayer to expand. However, this comes with an increase in computational costs.

### 3.2 Sorting of cardiolipins

Using scaffolding wall beads to apply a sinusoidal bilayer shape, we identified qualitative trends in curvature-driven sorting for a variety of different lipid types. First, our results confirm negative curvature preference for CDL as reported previously. In agreement with its high intrinsic curvature of -1.1 nm [19], we find strong enrichment of CDL in highly negatively curved membrane regions with *H* < - 0.2 nm^-1^. Matching the cone shape of CDL lipids with membrane curvature releases curvature frustration by optimizing the bending elasticity. However, we observed that various factors influence the magnitude of the observed sorting behavior: (i) Sorting of CDL varies strongly with the protonation state, i.e., highly charged CDL lipids have a lower affinity for negatively curved regions. Considering that in an active mitochondrion, the pH decreases from the proton source (electron transport chain) to the proton sink (ATP synthase), the pH gradient could lead to changes in the membrane shape, through dynamic cardiolipin (de)protonation [96]. (ii) The presence of divalent cations increases the curvature preference of CDL irrespective of the protonation state. This finding cannot be rationalized by the reduced head group repulsion alone. A recent study [73] suggests that Ca^2+^ ions not only induce short-range attractive forces between anionic lipids but also pronounced long-range lipid clustering that strongly correlates with negative curvature. Our findings are also in line with theoretical and experimental evidence that divalent cations can induce CDL clustering and the formation of buds in GUVs.[58, 97] More importantly, the influx of Ca^2+^ into the IMM triggers aggregation and immobilization of CDLs which reduces the interaction with respiratory chain complexes in mitochondria. Thus the clustering of CDL lipids in combination with enhanced curvature sorting might be essential for complex II to disintegrate which is a crucial step in cell death induction [98]. (iii) Furthermore, we observed that changing the lipid environment has a substantial impact on the sorting of CDL. The presence of POPE, another lipid with negative intrinsic curvature, causes competition for negatively curved regions [36], thereby reducing the sorting of CDL. On the other hand, in the presence of anionic lipids, more CDL lipids reside in negatively curved regions. According to X-ray diffraction studies of binary lipid mixtures, PS lipids experience positive intrinsic curvature [68], while the opposite was observed for PA lipids [99]. However, we only find enrichment of anionic lipids in positive curvature, thereby increasing the sorting behavior of CDL lipids. This apparent discrepancy might be explained by a possible difference in protonation. We modeled POPA in the dianionic state, while under physiological conditions POPA lipids are likely a mixture of mono- and di-anionic species, which leads to a reduced head group repulsion. Moreover, Coulomb interactions in the absence of long-range electrostatics in Martini 3 might not be sufficient to describe the interactions of lipids in highly charged bilayers. Further atomistic simulations are necessary to verify these findings. (iv) Finally, we saw that increasing the concentration of CDL reduces the preference for negative curvature following a reduction in CPP. However, at CDL concentrations above 20%, the sorting behavior can no longer be simply inferred from molecular shape considering that the packing of cone-shaped CDL lipids in a lamellar bilayer becomes increasingly difficult [38, 100].

### 3.3 Mixing entropy does not prevent lipid sorting

Despite numerous evidence of CDL sorting to negatively curved membrane regions [15, 19, 35, 36], other theoretical works [13, 23, 25, 66] suggest that the lipid mixing entropy dominates the lateral distribution of lipids. The authors claim, that strong lipid-protein or lipid-lipid interactions are required to enable efficient sorting relevant in biological systems. Using a nearest neighbor approximation to calculate the conditional mixing entropy, we find that sorting of CDL lipids in curved bilayers does not reduce the local mixing compared to the flat reference system. We hypothesize that lipid mixing in flat membranes could also be non-ideal due to mismatches in lipid shapes. On the other hand, an asymmetric lipid density in curved membranes can relieve this packing stress thereby enhancing the local mixing in curved regions. Given the conditional entropy as a metric for lipid mixing, these results show that arguments on the mixing of lipids in curved systems are more complex than generally assumed. The global reduction in entropy due to curvature-driven sorting could be less severe considering the increased local mixing. Furthermore, free energy calculations of moving a single CDL molecule across a curved bilayer show an energy difference of 3-4 *k*_*B*_*T* between negatively and positively curved regions depending on the CDL protonation state. Taken together, our results indicate that sorting of CDL in simple binary mixtures is not limited by the mixing entropy and does not require cooperative lipid-lipid interactions. The cone-shaped character is sufficient to drive the sorting of CDL lipids to negatively curved regions. Although this might only be true for membranes with considerable curvature gradients, the consequences could still be important in the context of biological processes. For example, the fusion of the IMM is regulated by the interaction between CDL and GTPase optic atrophy 1 (OPA1). Ban *et al*. [101] discovered that in the presence of low CDL concentrations IMMs are tethered by homotypic OPA1 which might stabilize the cristae morphology. Interestingly, OPA1 becomes fusogenic when CDL-rich domains are present in the opposing IMM, indicating that CDL’s preference for negative curvature might play a crucial role in the stalk formation between two IMMs. This result is in line with a recent computational study [102], showing that the free energies of stalk formation are significantly lower for cone-shaped lipids because their shape is most compatible with the large negative curvatures at the stalk rim.

### 3.4 Sorting induced by different lipid tails and head groups

Due to its unique four-tail structure, sorting of CDL is the most prominent example of how lipid shape can induce lateral segregation. However, we identified several other lipid types that experience curvature preferences to varying extents. In general, the sorting behavior of phospholipids is heavily affected by (i) the tail length, (ii) the degree of tail saturation, and (iii) head group type. Subsequently, we will discuss those three effects in more detail and highlight the implications on relevant biological processes.

i. We observed a strong correlation between lipids with short hydrocarbon tails and positively curved regions, although short-chain lipids are considerably cone-shaped according to our shape analysis. With increasing tail length, the preference for positive curvature decreases. However, no enrichment in negatively curved membrane regions could be observed despite computational and experimental evidence that most PC lipids have zero or slightly negative spontaneous curvatures. [21, 103, 104] While this result seems counterintuitive and contradicts the shape concept, it underlines the importance of lipid packing. Filling the gaps between lipids in positively curvature regions reduces the packing defects and thereby the overall bending rigidity of the bilayer [105]. Interestingly, a previous study employing all-atom simulations [106] reported that the addition of cone-shaped lipids actively drives the formation of membrane defects that resemble those induced by positive curvature. Considering the existence of various regulatory pathways [107, 108] that promote the formation of cone-shaped lipids by shortening lipid tails or decreasing the degree of tail saturation, the introduction of packing defects and thus positive curvature might be an active means to control the shape and function of cellular organelles.
ii. Analogously, highly unsaturated lipids show strong enrichment in positive curvature as the degree of saturation is tightly linked to the lipid chain length. This means that for a given number of tail atoms, unsaturated tails tend to be shorter than saturated ones. Simulations of small liposomes and membrane tethers have shown that polyunsaturated tails in the positively curved outer monolayer have the propensity to fold back toward the interface [109], thereby filling the defects and releasing packing stress in the bilayer center. Recently, tether-pulling simulations on asymmetric membranes provided further evidence that polyunsaturated lipids are more adaptable to positive curvature than saturated ones [110, 111]. Taken together, these results suggest that the presence of polyunsaturated lipids increases the membrane deformability, which can rationalize the abundance of polyunsaturated lipids in synaptic vesicles [112] and in the cytosolic leaflet of the plasma membrane [113].
iii. We found good agreement between molecular shape and observed curvature preference for different types of lipids head groups. As a consequence of its large spontaneous curvature of -1/3 nm^-1^ [67], zwitterionic POPE shows visible enrichment in negative curvature. To a lesser extent, we find enrichment of mono-anionic lipids like POPS in negative curvature. Despite electrostatic repulsion between the head groups, computational and experimental studies [72, 73] have reported small negative spontaneous curvatures for PS lipids. Surprisingly, di-anionic POPA shows moderate enrichment in positive curvature. However, reducing the head group charge of POPA to -1 leads to sorting of POPA to negative curvature, similar to POPS and POPG. This result highlights the importance of pH and its implications on the curvature preference of lipids. Maer *et al*. [114] recently demonstrated how the lifetime and conductance of gramicidin channels are regulated by the spontaneous curvature of DOPS and DOPE lipids. Reducing the pH from 7 to 3 led to a decrease in the head group charge of DOPS and thus a more negative spontaneous curvature, which decreased the channel lifetime. Conversely, the authors reported an increase in channel lifetime for DOPE-containing membranes, when reducing the pH to 0 and or increasing it to 9. At both pH levels, charged head groups prevail, thereby reducing the negative spontaneous curvature. Considering that similar effects have been observed for other classes of transmembrane proteins, e.g., changes in pH alter the gating properties of alamethicin channels in PS-containing bilayers [115, 116], this behavior likely originates from general bilayer properties rather than specific lipid-protein interactions.

### 3.5 Non-intuitive sorting in domain forming mixtures

While curvature preference and lipid packing govern the lateral distribution of simple binary lipid mixtures in curved bilayers, we also identified several factors that can lead to non-intuitive lipid sorting in more complex mixtures including lipid phase separation, interleaflet coupling, and line tension. Unfortunately, the contribution of these effects cannot be disentangled easily as they are intricately linked to each other. Nonetheless, the balance of forces can be shifted under certain conditions, which will be discussed below.

Lipid mixtures with significant differences in their miscibility tend to form co-existing nanodomains or, usually in the presence of CHOL, *L*_*o*_ and *L*_*d*_ phases. In both cases, the presence of interleaflet coupling leads to the registration of domains, which impedes the curvature-driven sorting of lipids. The larger the difference in tail length and saturation between the domains, the stronger is the phase separation and thus the domain registration [83, 117]. However, if the hydrophobic length mismatch between the lipid types is too large (e.g., di-16:3 vs di-20:0), no interleaflet coupling was observed. Instead, the release of packing defects directs polyunsaturated lipids to positively curved bilayer regions. Although the role of lipid tail interdigitation as one of the main coupling mechanisms has been heatedly debated [81], this result suggests that both lipid phases require at least a certain proximity to the bilayer midplane to be coupled effectively and outcompete curvature and packing preferences. At the domain boundaries, line tension arises as a consequence of the hydrophobic mismatch, facilitating the formation of continuous domains. In the strongly phase-separated system, line tension can push the stiffer *L*_*o*_ phase into less preferred, highly curved regions at the expense of membrane bending energy. By increasing the *L*_*d*_*/L*_*o*_ ratio, line tension can be decreased, which leads to the formation of non-continuous *L*_*d*_ phases that locate at highly curved regions. Accordingly, sorting of the *L*_*o*_ phase to zero curvature is promoted. An alternative way to restore curvature matching in phase-separated systems is to reduce the line tension by decreasing the hydrophobic mismatch between the *L*_*d*_ and *L*_*o*_ phases. Consequently, the level of phase separation is reduced, which alleviates the interleaflet coupling, leading to the enrichment of the unsaturated *L*_*d*_ phase in positively curved regions of both leaflets.

Interestingly, a recent experimental study [118] revealed that membrane curvature can also mediate interleaflet coupling in asymmetric LUVs composed of POPC and POPE. When POPC was enriched in the outer leaflet and POPE in the inner leaflet, cooperative melting and similar packing of both lipid types in the two leaflets occurred. When reversing the leaflet composition, i.e., enriching the outer leaflet with POPE and the inner with POPC, the leaflets melted independently and the coupling disappeared. The authors concluded that the energetically preferred position of POPE is in the inner leaflet, consistent with its cone shape and natural occurrence in the cytosolic leaflet of the plasma membrane, which induces curvature-mediated coupling between the asymmetric leaflets. Considering that both curvature [111] and lipid phase separation [119] can drive the lateral segregation of proteins, the interplay between the two could be fundamental for the localization of proteins in cellular processes.

### 3.6 Conclusions

To conclude, we demonstrated that using artificial wall beads is a simple yet effective way to reshape lipid bilayers. Arbitrary curvature gradients can be precisely controlled without affecting the underlying lipid organization and dynamics. When applying curvature to bilayers, lipids experience curvature-driven sorting qualitatively matching their intrinsic shape. The more their shape deviates from a cylindrical shape, the stronger is the sorting effect as shown for different protonation states of cardiolipin. Surprisingly, the presented data suggest that the sorting of cardiolipin does not oppose lipid mixing entropy and does not require cooperative lipid-lipid or lipid-protein interactions. Considering the large intrinsic curvature of cardiolipin and that its concentration is actively regulated in the mitochondria, curvature-driven sorting of cardiolipin could be vital for processes like membrane fusion of the OMM and IMM. For various other lipid types, we found that the sorting preference is largely determined by the headgroup type. However, the packing preference of lipid tails, which depends on the effective tail length, can strongly alter the observed curvature sorting. Especially polyunsaturated lipids show highly promiscuous sorting behavior depending on their environment. In more complex lipid mixtures, non-intuitive sorting can arise from a delicate balance between intrinsic curvature preference and other forces, e.g., lipid phase separation, line tension, and interleaflet coupling. Despite current beliefs that curvature-driven sorting is inferior to those other forces, we identified several scenarios where lipid curvature or packing preferences were key in determining the lateral lipid distribution. The remarkable ability to alter the lateral organization of lipid mixtures indicates that curvature-driven sorting might be essential in various cellular pathways.

In addition, extensive knowledge about the lateral distribution of lipids in membranes with complex compositions and curvature gradients can accelerate the simulation of realistic membranes. By providing a vast training set for AI-based structure builders, close-to-equilibrium structures can be generated which will pave the way to simulating entire organelles and ultimately a whole cell at near atomistic resolution.

## 4. Methods

All MD simulations were carried out using different versions of Gromacs (2018-2022) [120] with the Martini 3 force field [121]. Initially, parameters were taken from the beta-release of Martini 3, later using the final release with no significant differences in the results except for systems involving Ca^2+^ ions. In those cases, the final Martini 3 parameters behaved more realistically, and only those results are included.

Flat and curved membranes were set up using version 1.1 of TS2CG [88]. To maintain the desired curvature, wall beads were placed above the lipid head group region of both leaflets. Wall beads interact repulsively with lipid tail beads (type C1 and C4h for saturated and unsaturated fragments, respectively) and they were frozen over the course of the simulation to keep the membrane in a sine shape.

All unbiased simulations of curved membranes were performed for 5 µs (binary systems) or 10 µs (ternary systems) and in replicates of three. PMFs were computed by umbrella sampling, pulling a single CDL lipid across a curved bilayer. For each CDL protonation state, 260 windows were extracted and simulated for 1 µs.

Further details on the setup of flat and curved membranes, simulation parameters, and analysis are provided in the Supplementary Information including a table listing all simulations performed (Supplementary Table 2).

## Acknowledgment

The authors thank A.H. de Vries for the discussion and feedback regarding umbrella sampling simulations of curved membranes. We would like to thank the Center for Information Technology of the University of Groningen for their support and for providing access to the Peregrine/Hábrók high-performance computing cluster and SURFsara for providing access to Snellius. This research is supported by the “BaSyC—Building a Synthetic Cell” Gravitation grant (024.003.019) of the Netherlands Ministry of Education, Culture, and Science (OCW) and the Netherlands Organization for Scientific Research (NWO).

