## Supplementary Information for "Curvature-induced lipid sorting beyond the critical packing parameter"

### Supplementary Tables

**Supplementary Table 1 Area per lipid (APL), tail volume ($v_0$), tail length ($l_c$), and resulting critical packing parameter (CPP) of Martini models for CDL0, CDL1, and CDL2.** All values are computed from flat bilayer simulations for a 9:1 mixture of POPC/CDL in the presence of 150 mmol NaCl or 50 mmol $CaCl_2$ (plus counterions). The size of the headgroup beads is the same for all the CDL protonation states, only the charge changes. No difference between Martini 3 beta and Martini 3 final could be observed in the presence of sodium ions. CDL values in the presence of calcium ions come from simulations using Martini 3 final parameters. Errors indicate standard deviation during the last 200 ns of a single simulation.

| | Lipid | APL [$nm^2$] | $v_0$ [$nm^3$] | $l_c$ [nm] | CPP |
|---|---|---|---|---|---|
| $Na^+$ | CDL0 | 0.94 ± 0.03 | 1.51 ± 0.01 | 1.17 ± 0.10 | 1.40 ± 0.14 |
| | CDL1 | 1.02 ± 0.03 | 1.51 ± 0.01 | 1.18 ± 0.10 | 1.26 ± 0.11 |
| | CDL2 | 1.13 ± 0.03 | 1.51 ± 0.01 | 1.20 ± 0.10 | 1.12 ± 0.10 |
| $Ca^{2+}$ | CDL0 | 0.93 ± 0.03 | 1.51 ± 0.01 | 1.16 ± 0.10 | 1.40 ± 0.14 |
| | CDL1 | 1.01 ± 0.03 | 1.51 ± 0.01 | 1.18 ± 0.10 | 1.27 ± 0.12 |
| | CDL2 | 1.09 ± 0.03 | 1.51 ± 0.01 | 1.21 ± 0.10 | 1.15 ± 0.10 |

**Supplementary Table 2 List of simulations.** All systems with a curvature amplitude of 0 denote flat reference systems simulated in the absence of confining wall beads (except stated otherwise). The initial system size of flat reference systems was set to 15 x 10 x 10 $nm^3$ containing between 175 to 215 lipids per leaflet (depending on the membrane composition) and 6600 to 7800 water molecules. All systems with a curvature amplitude of 4 denote curved systems simulated in the presence of confining wall beads. The initial system size of binary curved systems was set to 20 x 10 x 15 $nm^3$ containing between 266 to 364 lipids per leaflet (depending on the membrane composition) and 15,000 to 17,000 water molecules. All systems (flat and curved) were neutralized and 150 mmol of NaCl was added (or 50 mmol of $CaCl_2$).

[a] To compare the conditional entropy of mixing of flat and curved systems, they need to have the same number of lipids. Therefore, another set of flat simulations with a larger box size was performed.

[b] Umbrella sampling simulations of pulling a single cardiolipin molecule across the entire length of the curved membrane.

[c] To allow for proper phase separation, larger box sizes of 20 x 20 x 15 $nm^3$ were used. For curved simulations, one simulation with a random starting configuration and two simulations with biased starting configurations were performed.

| Membrane composition | Lipid ratio | Curvature ampl. [nm] | Time [µs] | Replicas | Special conditions |
|---|---|---|---|---|---|
| ***Influence of wall beads on flat membrane (Fig. 1)*** | | | | | |
| POPC | | 0 | 1 | 1 | |
| POPC | | 0 | 1 | 1 | with wall beads |
| ***Sorting of neutral cardiolipin (CDL0) in POPC membrane (Fig. 2)*** | | | | | |
| POPC/CDL0 | 9/1 | 0 | 5 | 1 | |
| POPC/CDL0 | 9/1 | 4 | 5 | 3 | |
| ***Sorting of cardiolipin with different protonation states in POPC membrane (Fig. 3)*** | | | | | |
| POPC/CDL0 | 9/1 | 0 | 5 | 1 | |
| POPC/CDL0 | 9/1 | 4 | 5 | 3 | |
| POPC/CDL1 | 9/1 | 0 | 5 | 1 | |
| POPC/CDL1 | 9/1 | 4 | 5 | 3 | |
| POPC/CDL2 | 9/1 | 0 | 5 | 1 | |
| POPC/CDL2 | 9/1 | 4 | 5 | 3 | |
| ***Effect of ions on sorting behavior of cardiolipin (Fig. 4)*** | | | | | |
| POPC/CDL0 | 9/1 | 0 | 0.5 | 1 | |
| POPC/CDL0 | 9/1 | 4 | 5 | 3 | |
| POPC/CDL1 | 9/1 | 0 | 0.5 | 1 | |
| POPC/CDL1 | 9/1 | 4 | 5 | 3 | |
| POPC/CDL2 | 9/1 | 0 | 0.5 | 1 | |
| POPC/CDL2 | 9/1 | 4 | 5 | 3 | |
| POPC/CDL0 | 9/1 | 0 | 0.5 | 1 | $CaCl_2$ |
| POPC/CDL0 | 9/1 | 4 | 5 | 3 | $CaCl_2$ |
| POPC/CDL1 | 9/1 | 0 | 0.5 | 1 | $CaCl_2$ |
| POPC/CDL1 | 9/1 | 4 | 5 | 3 | $CaCl_2$ |
| POPC/CDL2 | 9/1 | 0 | 0.5 | 1 | $CaCl_2$ |

| | | | | | |
|---|---|---|---|---|---|
| POPC/CDL2 | 9/1 | 4 | 5 | 3 | $CaCl_2$ |
| ***Sorting of cardiolipin in various bulk lipids (Fig. 5)*** | | | | | |
| POPA/CDL0 | 9/1 | 0 | 0.5 | 1 | |
| POPA/CDL0 | 9/1 | 4 | 5 | 3 | |
| POPA/CDL1 | 9/1 | 0 | 0.5 | 1 | |
| POPA/CDL1 | 9/1 | 4 | 5 | 3 | |
| POPA/CDL2 | 9/1 | 0 | 0.5 | 1 | |
| POPA/CDL2 | 9/1 | 4 | 5 | 3 | |
| POSM/CDL0 | 9/1 | 0 | 0.5 | 1 | |
| POSM/CDL0 | 9/1 | 4 | 5 | 3 | |
| POSM/CDL1 | 9/1 | 0 | 0.5 | 1 | |
| POSM/CDL1 | 9/1 | 4 | 5 | 3 | |
| POSM/CDL2 | 9/1 | 0 | 0.5 | 1 | |
| POSM/CDL2 | 9/1 | 4 | 5 | 3 | |
| POPS/CDL0 | 9/1 | 0 | 0.5 | 1 | |
| POPS/CDL0 | 9/1 | 4 | 5 | 3 | |
| POPS/CDL1 | 9/1 | 0 | 0.5 | 1 | |
| POPS/CDL1 | 9/1 | 4 | 5 | 3 | |
| POPS/CDL2 | 9/1 | 0 | 0.5 | 1 | |
| POPS/CDL2 | 9/1 | 4 | 5 | 3 | |
| POPG/CDL0 | 9/1 | 0 | 0.5 | 1 | |
| POPG/CDL0 | 9/1 | 4 | 5 | 3 | |
| POPG/CDL1 | 9/1 | 0 | 0.5 | 1 | |
| POPG/CDL1 | 9/1 | 4 | 5 | 3 | |
| POPG/CDL2 | 9/1 | 0 | 0.5 | 1 | |
| POPG/CDL2 | 9/1 | 4 | 5 | 3 | |
| POPC/CDL0 | 9/1 | 0 | 0.5 | 1 | |
| POPC/CDL0 | 9/1 | 4 | 5 | 3 | |
| POPC/CDL1 | 9/1 | 0 | 0.5 | 1 | |
| POPC/CDL1 | 9/1 | 4 | 5 | 3 | |
| POPC/CDL2 | 9/1 | 0 | 0.5 | 1 | |
| POPC/CDL2 | 9/1 | 4 | 5 | 3 | |
| POPE/CDL0 | 9/1 | 0 | 0.5 | 1 | |
| POPE/CDL0 | 9/1 | 4 | 5 | 3 | |
| POPE/CDL1 | 9/1 | 0 | 0.5 | 1 | |
| POPE/CDL1 | 9/1 | 4 | 5 | 3 | |
| POPE/CDL2 | 9/1 | 0 | 0.5 | 1 | |
| POPE/CDL2 | 9/1 | 4 | 5 | 3 | |
| ***Effect of concentration on the sorting of cardiolipin in POPC (Fig. 6)*** | | | | | |
| POPC/CDL0 | 95/5 | 0 | 0.5 | 1 | |
| POPC/CDL0 | 95/5 | 4 | 5 | 3 | |
| POPC/CDL0 | 95/5 | 0 | 5 | 1 | [a] large system for entropy calc. |
| POPC/CDL0 | 90/10 | 0 | 0.5 | 1 | |
| POPC/CDL0 | 90/10 | 4 | 5 | 3 | |
| POPC/CDL0 | 90/10 | 0 | 5 | 1 | [a] large system for entropy calc. |
| POPC/CDL0 | 85/15 | 0 | 0.5 | 1 | |
| POPC/CDL0 | 85/15 | 4 | 5 | 3 | |
| POPC/CDL0 | 85/15 | 0 | 5 | 1 | [a] large system for entropy calc. |
| POPC/CDL0 | 80/20 | 0 | 0.5 | 1 | |
| POPC/CDL0 | 80/20 | 4 | 5 | 3 | |
| POPC/CDL0 | 80/20 | 0 | 5 | 1 | [a] large system for entropy calc. |
| POPC/CDL0 | 75/25 | 0 | 0.5 | 1 | |
| POPC/CDL0 | 75/25 | 4 | 5 | 3 | |
| POPC/CDL0 | 75/25 | 0 | 5 | 1 | [a] large system for entropy calc. |
| POPC/CDL0 | 70/30 | 0 | 0.5 | 1 | |
| POPC/CDL0 | 70/30 | 4 | 5 | 3 | |

| | | | | | |
|---|---|---|---|---|---|
| POPC/CDL0 | 70/30 | 0 | 5 | 1 | [a] large system for entropy calc. |
| POPC/CDL0 | 65/35 | 0 | 0.5 | 1 | |
| POPC/CDL0 | 65/35 | 4 | 5 | 3 | |
| POPC/CDL0 | 65/35 | 0 | 5 | 1 | [a] large system for entropy calc. |
| POPC/CDL0 | 60/40 | 0 | 0.5 | 1 | |
| POPC/CDL0 | 60/40 | 4 | 5 | 3 | |
| POPC/CDL0 | 60/40 | 0 | 5 | 1 | [a] large system for entropy calc. |
| POPC/CDL0 | 55/45 | 0 | 0.5 | 1 | |
| POPC/CDL0 | 55/45 | 4 | 5 | 3 | |
| POPC/CDL0 | 55/45 | 0 | 5 | 1 | [a] large system for entropy calc. |
| POPC/CDL0 | 50/50 | 0 | 0.5 | 1 | |
| POPC/CDL0 | 50/50 | 4 | 5 | 3 | |
| POPC/CDL0 | 50/50 | 0 | 5 | 1 | [a] large system for entropy calc. |
| ***Free energy of cardiolipin sorting (Fig 7)*** | | | | | |
| POPC/CDL0 | 1 CDL0 | 0 | 0.5 | 1 | |
| POPC/CDL0 | 1 CDL0 | 0 | 1 | 1 | [b] US: 256 windows |
| POPC/CDL1 | 1 CDL1 | 0 | 0.5 | 1 | |
| POPC/CDL1 | 1 CDL1 | 0 | 1 | 1 | [b] US: 256 windows |
| POPC/CDL2 | 1 CDL2 | 0 | 0.5 | 1 | |
| POPC/CDL2 | 1 CDL2 | 0 | 1 | 1 | [b] US: 256 windows |
| ***Effect of tail length, tail saturation and headgroup type on the sorting of other phospholipids (Fig. 8)*** | | | | | |
| DNPC/DTPC | 9/1 | 0 | 0.5 | 1 | |
| DNPC/DTPC | 9/1 | 4 | 5 | 3 | |
| DNPC/DLPC | 9/1 | 0 | 0.5 | 1 | |
| DNPC/DLPC | 9/1 | 4 | 5 | 3 | |
| DNPC/LPPC | 9/1 | 0 | 0.5 | 1 | |
| DNPC/LPPC | 9/1 | 4 | 5 | 3 | |
| DNPC/DPPC | 9/1 | 0 | 0.5 | 1 | |
| DNPC/DPPC | 9/1 | 4 | 5 | 3 | |
| DNPC/DBPC | 9/1 | 0 | 0.5 | 1 | |
| DNPC/DBPC | 9/1 | 4 | 5 | 3 | |
| DNPC/DXPC | 9/1 | 0 | 0.5 | 1 | |
| DNPC/DXPC | 9/1 | 4 | 5 | 3 | |
| DNPC/DFPC | 9/1 | 0 | 0.5 | 1 | |
| DNPC/DFPC | 9/1 | 4 | 5 | 3 | |
| DNPC/DIPC | 9/1 | 0 | 0.5 | 1 | |
| DNPC/DIPC | 9/1 | 4 | 5 | 3 | |
| DNPC/DOPC | 9/1 | 0 | 0.5 | 1 | |
| DNPC/DOPC | 9/1 | 4 | 5 | 3 | |
| DNPC/POPC | 9/1 | 0 | 0.5 | 1 | |
| DNPC/POPC | 9/1 | 4 | 5 | 3 | |
| DNPC/DPPC | 9/1 | 0 | 0.5 | 1 | |
| DNPC/DPPC | 9/1 | 4 | 5 | 3 | |
| POPC/POPA | 9/1 | 0 | 0.5 | 1 | |
| POPC/POPA | 9/1 | 4 | 5 | 3 | |
| POPC/POSM | 9/1 | 0 | 0.5 | 1 | |
| POPC/POSM | 9/1 | 4 | 5 | 3 | |
| POPC/POPS | 9/1 | 0 | 0.5 | 1 | |
| POPC/POPS | 9/1 | 4 | 5 | 3 | |
| POPC/POPG | 9/1 | 0 | 0.5 | 1 | |
| POPC/POPG | 9/1 | 4 | 5 | 3 | |
| POPC/POPE | 9/1 | 0 | 0.5 | 1 | |
| POPC/POPE | 9/1 | 4 | 5 | 3 | |
| ***Effect of nanodomain formation on sorting of DFPC (Fig. 9)*** | | | | | |
| DIPC/DFPC | 9/1 | 0 | 0.5 | 1 | |
| DIPC/DFPC | 9/1 | 4 | 5 | 3 | |

| | | | | | |
|---|---|---|---|---|---|
| DOPC/DFPC | 9/1 | 0 | 0.5 | 1 | |
| DOPC/DFPC | 9/1 | 4 | 5 | 3 | |
| POPC/DFPC | 9/1 | 0 | 0.5 | 1 | |
| POPC/DFPC | 9/1 | 4 | 5 | 3 | |
| DPPC/DFPC | 9/1 | 0 | 0.5 | 1 | |
| DPPC/DFPC | 9/1 | 4 | 5 | 3 | |
| DTPC/DFPC | 9/1 | 0 | 0.5 | 1 | |
| DTPC/DFPC | 9/1 | 4 | 5 | 3 | |
| DLPC/DFPC | 9/1 | 0 | 0.5 | 1 | |
| DLPC/DFPC | 9/1 | 4 | 5 | 3 | |
| LPPC/DFPC | 9/1 | 0 | 0.5 | 1 | |
| LPPC/DFPC | 9/1 | 4 | 5 | 3 | |
| DBPC/DFPC | 9/1 | 0 | 0.5 | 1 | |
| DBPC/DFPC | 9/1 | 4 | 5 | 3 | |
| DXPC/DFPC | 9/1 | 0 | 0.5 | 1 | |
| DXPC/DFPC | 9/1 | 4 | 5 | 3 | |
| ***Lipid sorting in ternary mixtures undergoing phase separation (Fig. 10)*** | | | | | |
| DPPC/CHOL/DFPC | 35/35/30 | 0 | 10 | 1 | [c] |
| DPPC/CHOL/DFPC | 35/35/30 | 4 | 10 | 3 | [c] including 2 biased starting config. |
| DPPC/CHOL/DIPC | 35/35/30 | 0 | 10 | 1 | [c] |
| DPPC/CHOL/DIPC | 35/35/30 | 4 | 10 | 3 | [c] including 2 biased starting config. |
| DPPC/CHOL/DFPC | 50/40/10 | 4 | 10 | 1 | [c]; Suppl. Fig. 11 |
| ***Sorting of cholesterol (Suppl. Fig. 12)*** | | | | | |
| POPC/CHOL | 9/1 | 0 | 0.5 | 1 | |
| POPC/CHOL | 9/1 | 0 | 5 | 1 | |

### Supplementary Figures

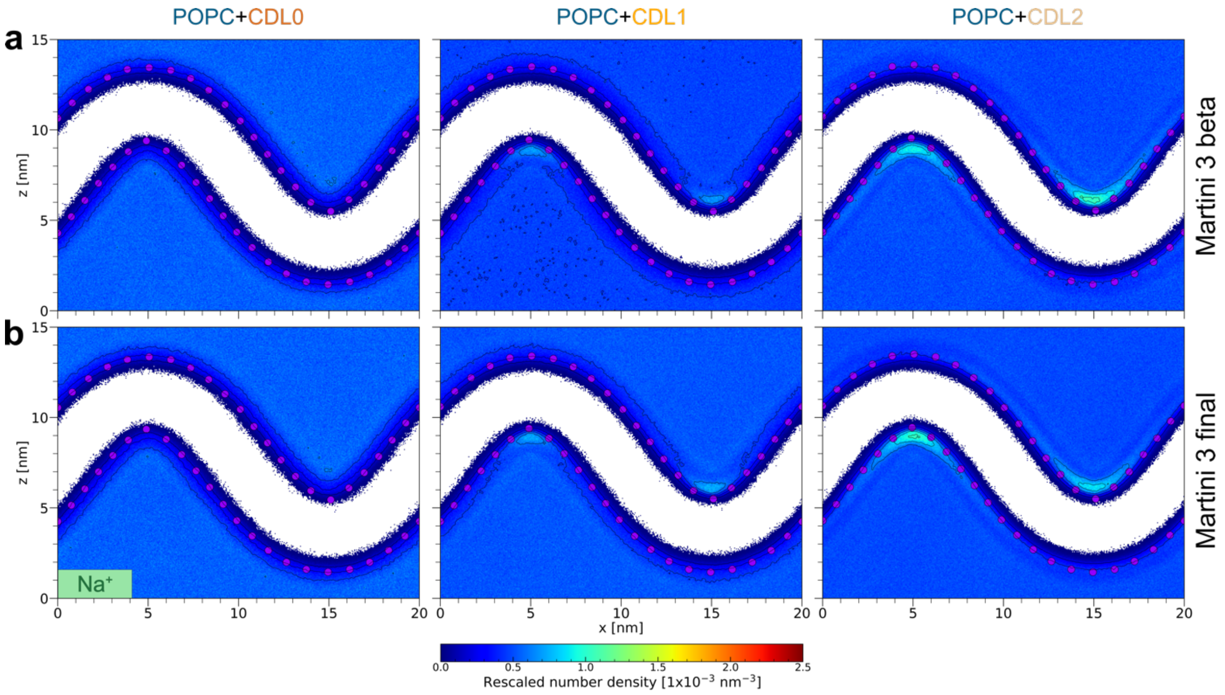


**Supplementary Figure 1 $Na^+$ distribution around CDL containing curved bilayers.** The 2d number density of $Na^+$ using (a) Martini 3 beta and (b) Martini 3 final parameters for neutral CDL0 (left), anionic CDL1 (middle), and double anionic CDl2 (right) is computed as an average over the last 2 µs of 3 replicates.

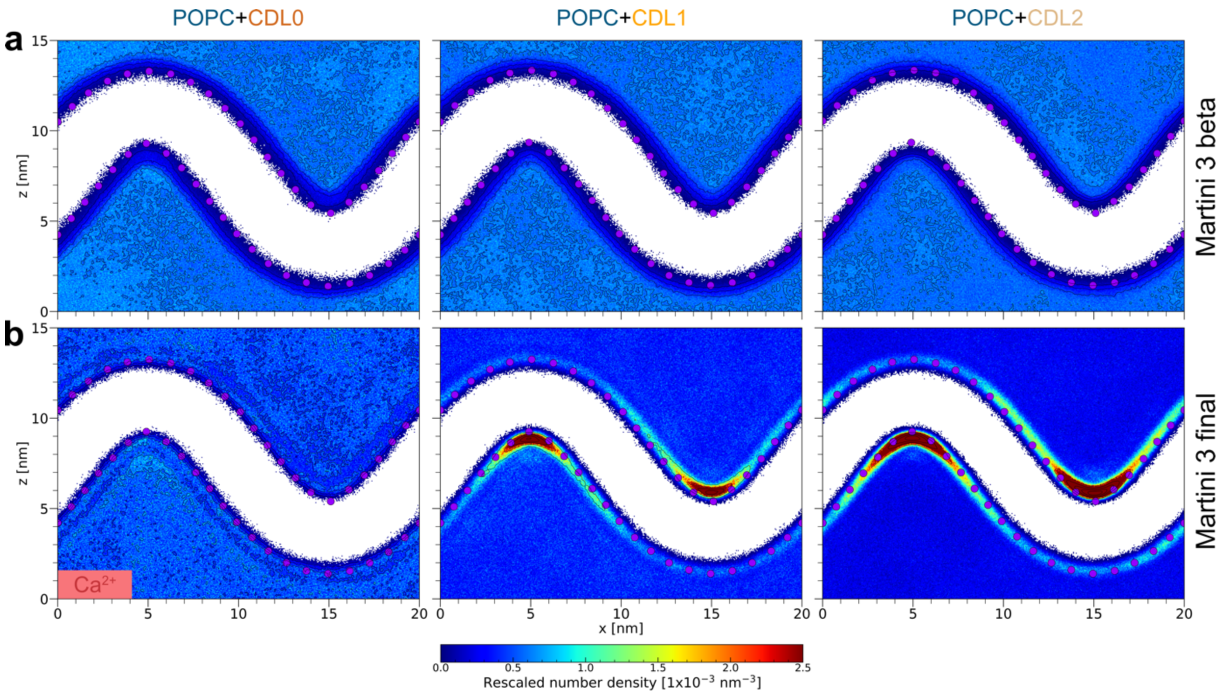


**Supplementary Figure 2 $Ca^{2+}$ distribution around CDL containing curved bilayers.** The 2d number density of $Ca^{2+}$ using (a) Martini 3 beta and (b) Martini 3 final parameters for neutral CDL0 (left), anionic CDL1 (middle), and double anionic CDl2 (right) is computed as an average over the last 2 µs of 3 replicates.

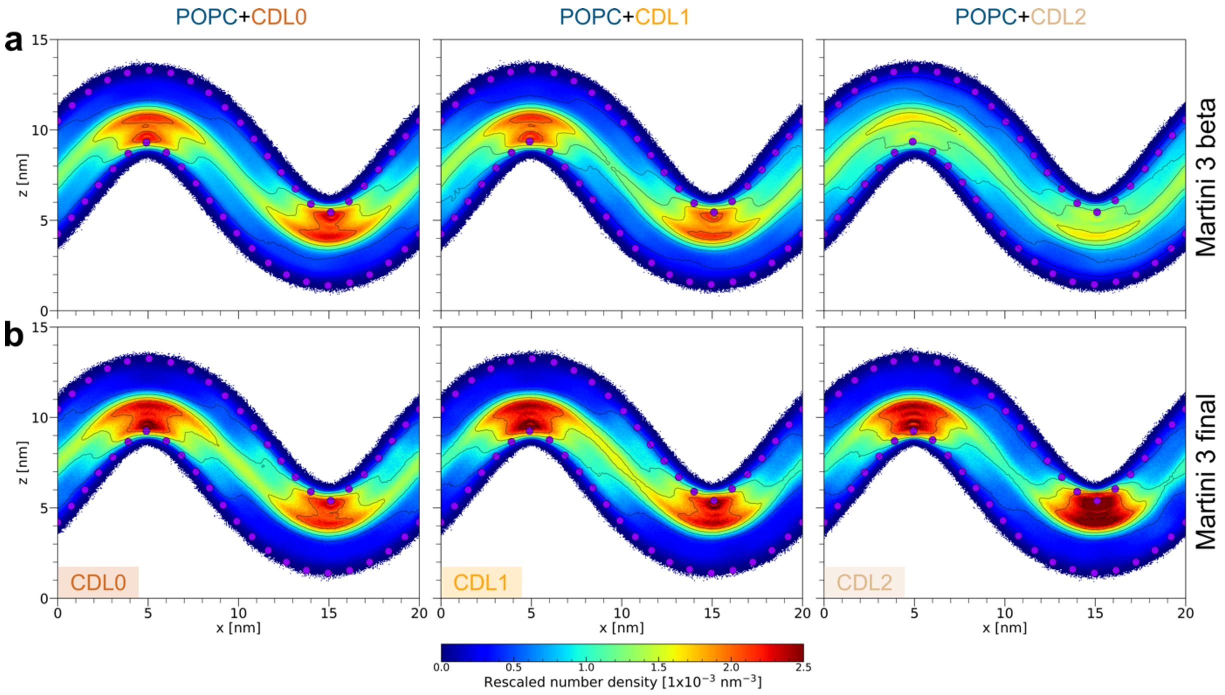


**Supplementary Figure 3 Distribution of CDL lipids in curved POPC membrane in the presence of $Ca^{2+}$.** The 2d number density of CDL using (a) Martini 3 beta and (b) Martini 3 final parameters for neutral CDL0 (left), anionic CDL1 (middle), and double anionic CDL2 (right) is computed as an average over the last 2 µs of 3 replicates.

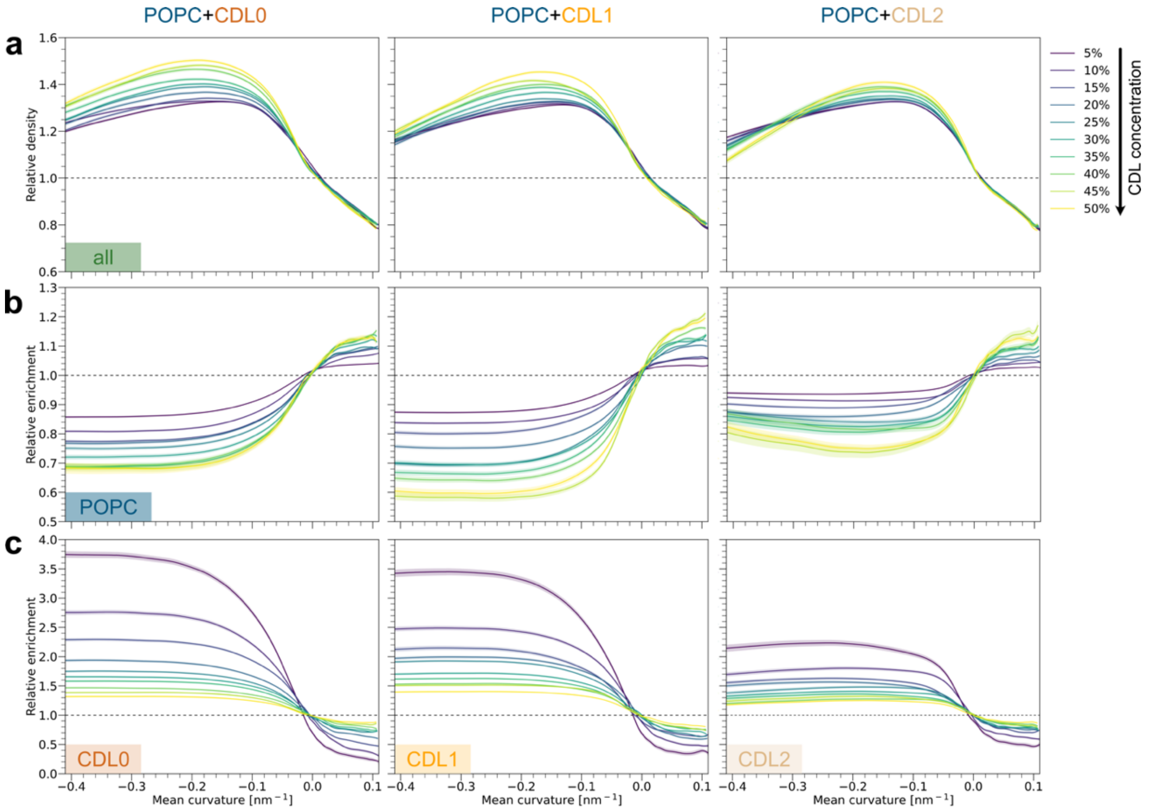


**Supplementary Figure 4 Distribution of CDL and POPC lipids in different curvatures.** (a) Relative density and (b and c) enrichment plots for the three different CDL protonation states (from left to right) at 10 different CDL concentrations.

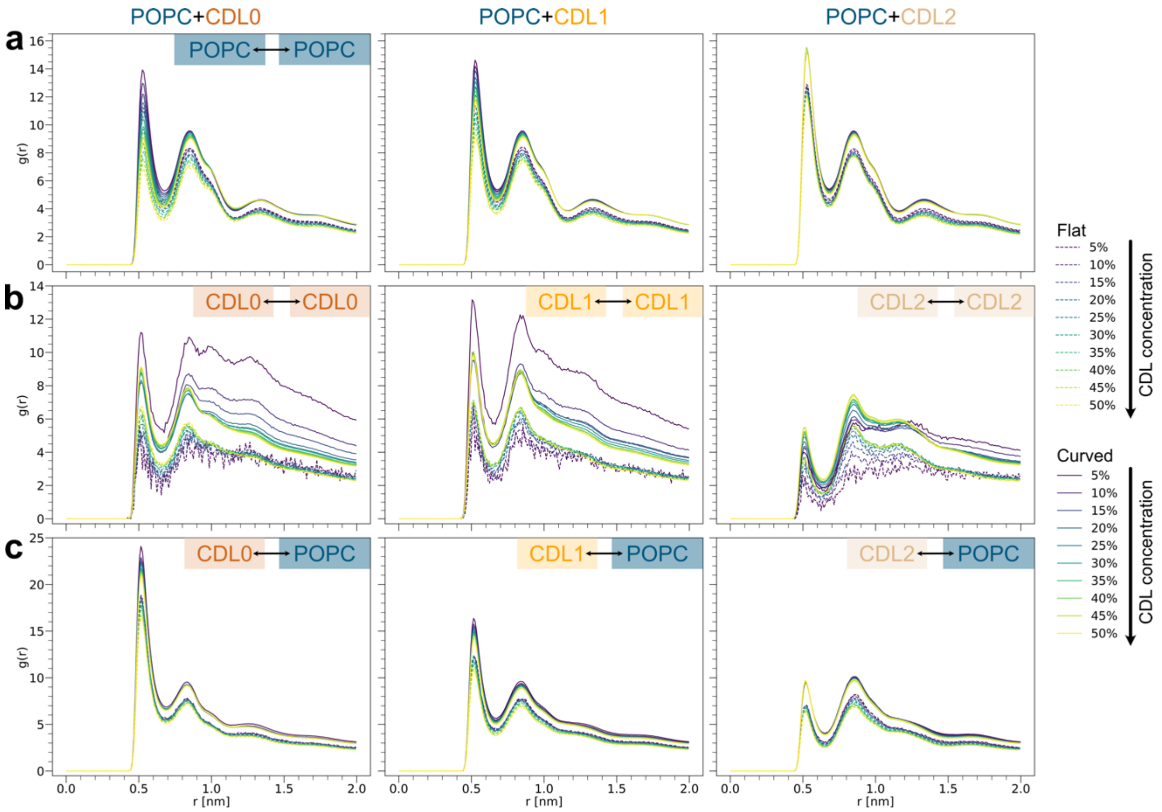


**Supplementary Figure 5 POPC/CDL lipid shells in flat and curved membranes.** Lipid-lipid RDFs for (a) POPC, (b) CDL-CDL, and (c) CDL-POPC for the three different CDL protonation states (from left to right) at 10 different CDL concentrations. RDFs are computed with respect to the central GL0 bead for CDL lipids and the PO4 bead for POPC lipids. All data referred to as ‘Flat’ (dashed lines) is computed from flat bilayer simulations with the same lipid ratios as ‘Curved’ systems (solid lines).

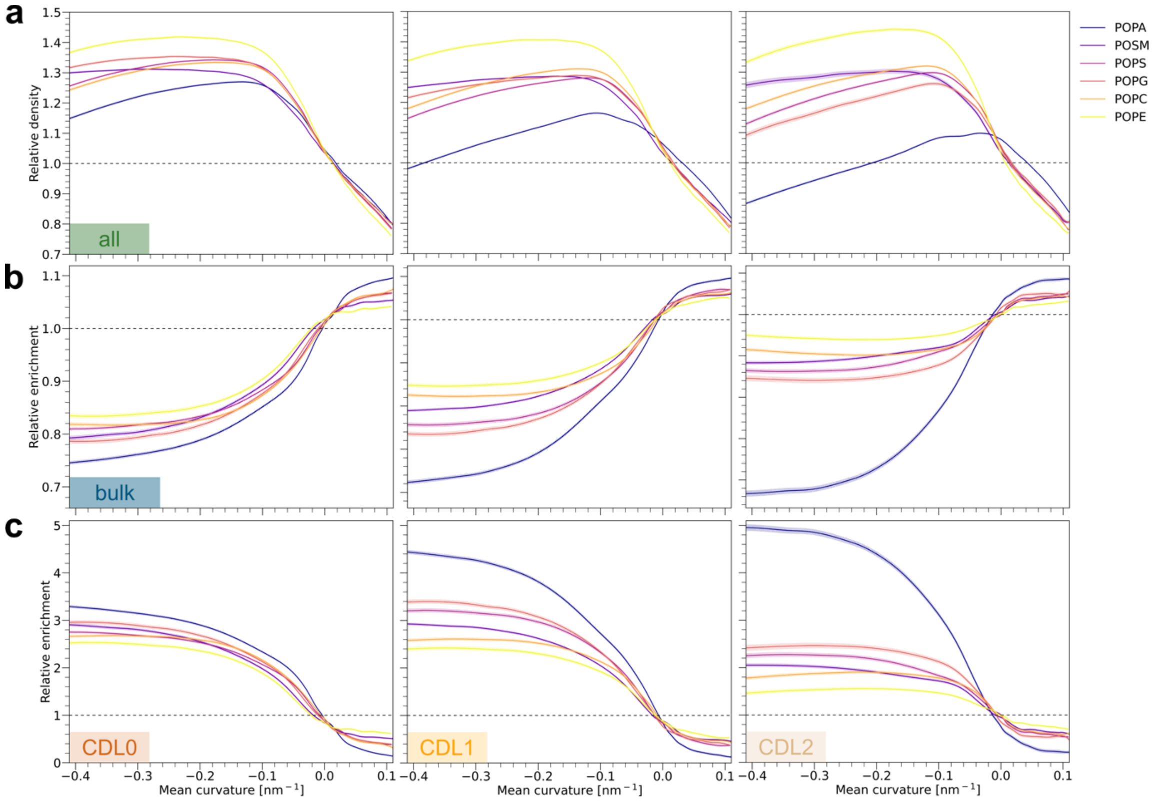


**Supplementary Figure 6 Distribution of CDL and bulk lipids in different curvatures.** (a) Relative density and (b and c) enrichment plots for the three different CDL protonation states (from left to right) in POPA, POSM, POPS, POPG, POPC, and POPE.

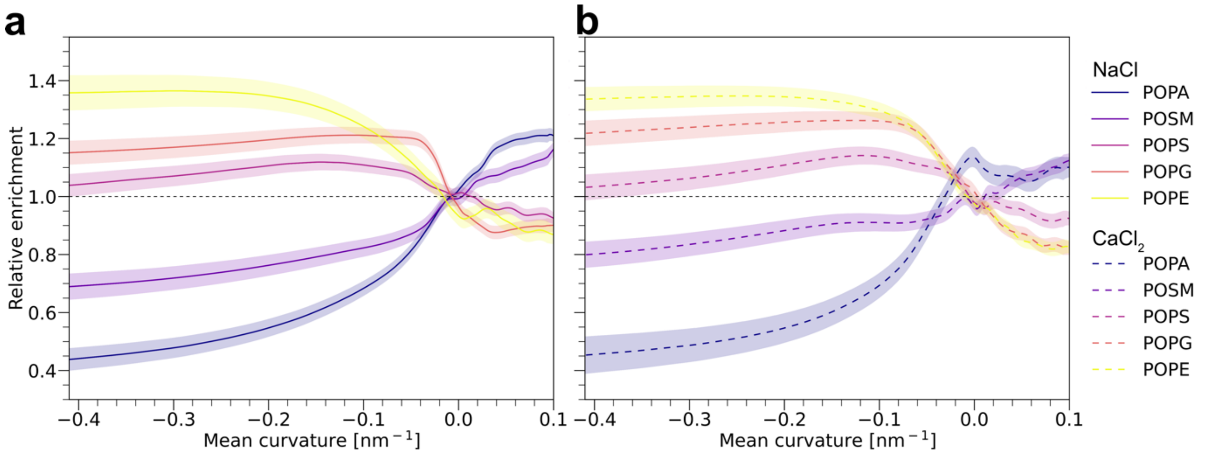


**Supplementary Figure 7 Effect of $Ca^{2+}$ on the enrichment of lipids with different head group types in POPC.** Relative enrichment of POPA, POSM, POPS, POPG, and POPE in POPC in the presence of (a) 150 mmol NaCl and (b) 50 mmol $CaCl_2$. The presence of $Ca^{2+}$ leads to minor enrichment of mono-anionic lipids (POPS and POPG) in negative curvature, while di-anionic POPA becomes enriched in zero-curvature regions. All data was generated using the final Martini 3 parameters.

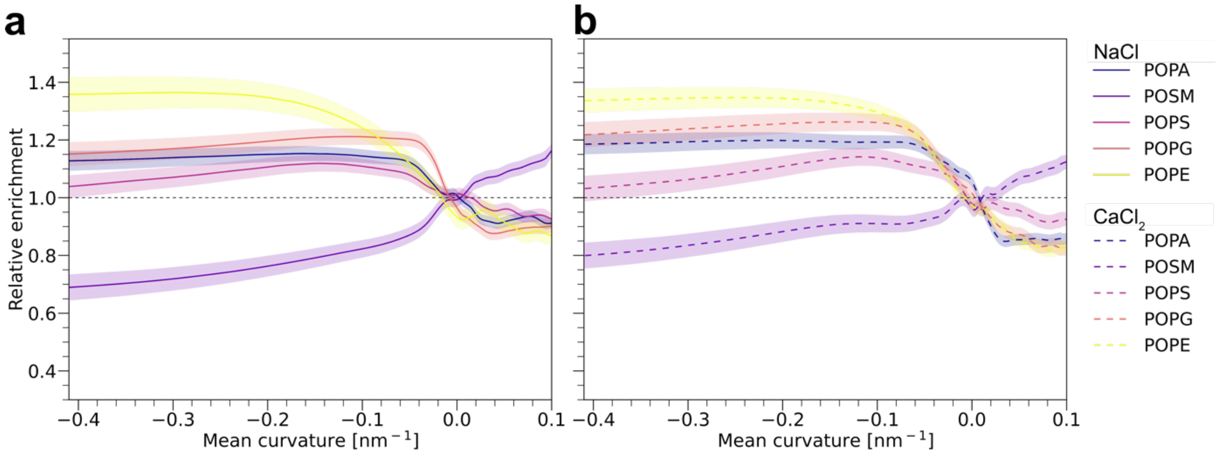


**Supplementary Figure 8 Effect of reduction in anionic charge on the sorting behavior of POPA.** Relative enrichment of POPA, POSM, POPS, POPG, and POPE in POPC in the presence of (a) 150 mmol NaCl and (b) 50 mmol $CaCl_2$. The presence of $Ca^{2+}$ leads to minor enrichment of mono-anionic POPA in negative curvature, comparable to POPS and POPG. All data was generated using the final Martini 3 parameters.

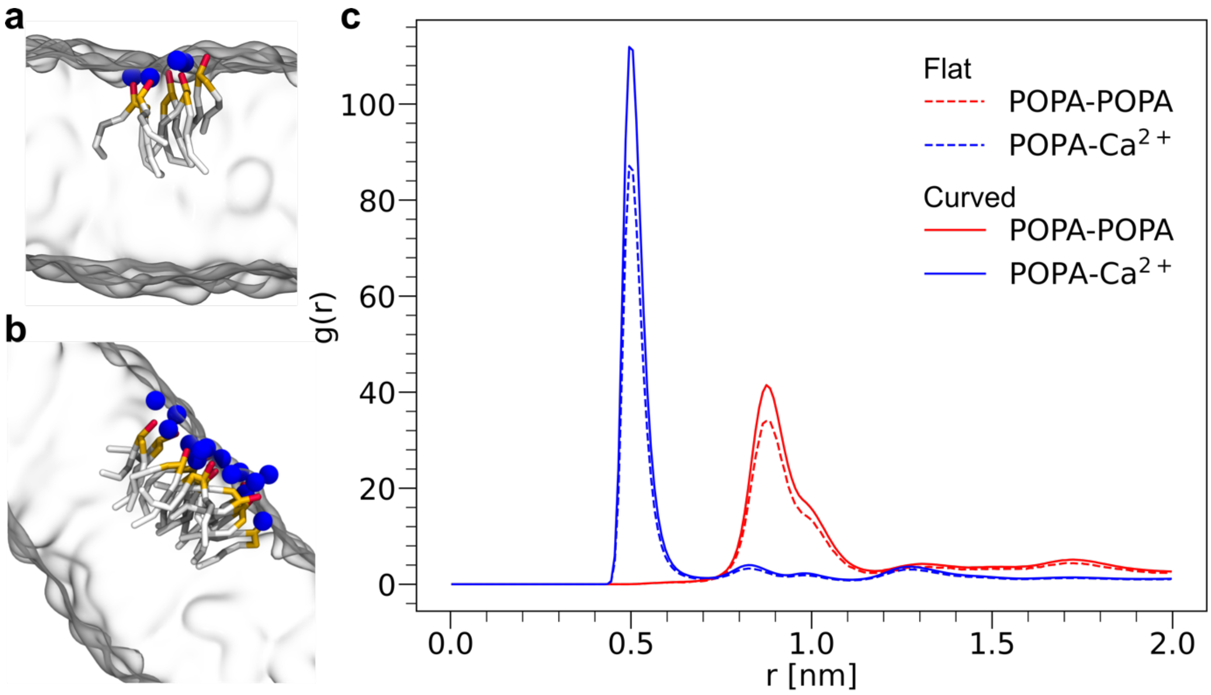


**Supplementary Figure 9 POPA lipid clusters in the presence of $Ca^{2+}$.** Representative snapshots of POPA cluster in (a) flat and (b) curved bilayer in a mixture with POPC (indicated as transparent surface) and (c) corresponding POPA-POPA and POPA-$Ca^{2+}$ All data referred to as 'Flat' (dashed lines) are computed from flat bilayer simulations with the same lipid ratio as 'Curved' systems (solid lines).

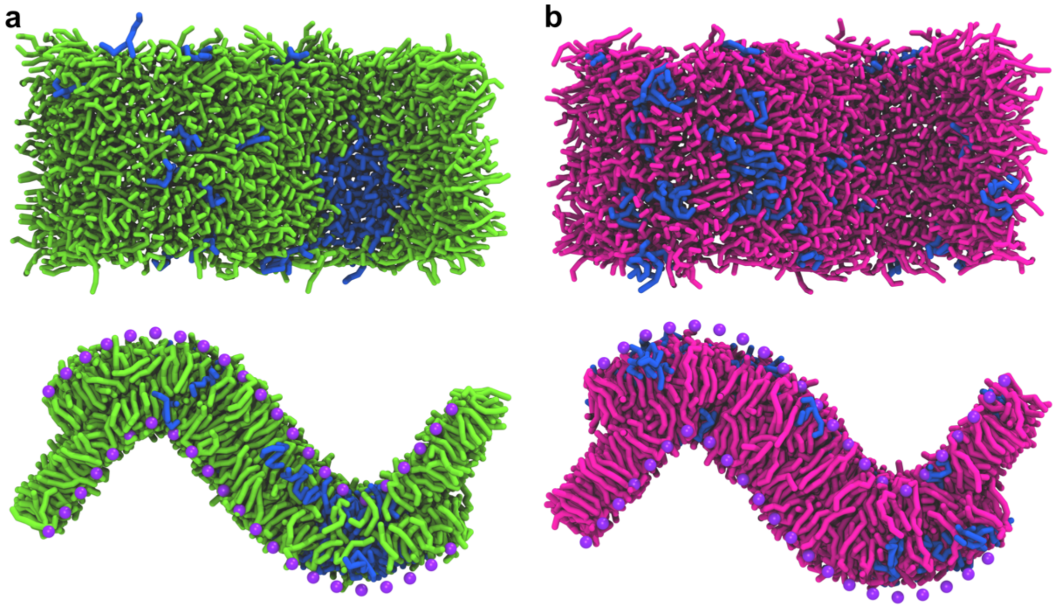


**Supplementary Figure 10 Snapshots of DFPC (blue) in binary mixtures with (a) DPPC (green) and (b) DBPC (magenta).** Both systems show visible formation of DFPC-rich nanodomains but remain in the fluid state.

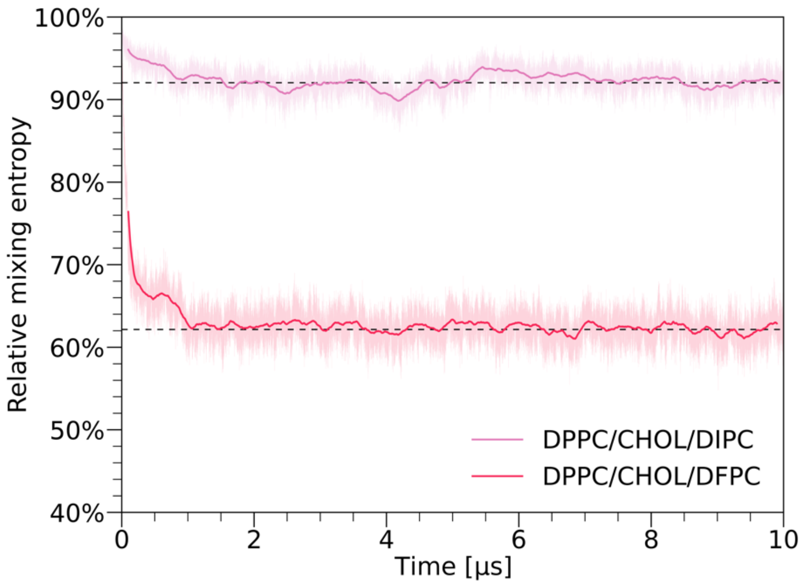


**Supplementary Figure 11 Relative conditional mixing entropy for weakly (DPPC/CHOL/DIPC) and strongly (DFPC/CHOL/DPPC) phase-separated systems.** Both simulations started with a random lipid distribution but showed quick (~ 1 µs) phase separation.

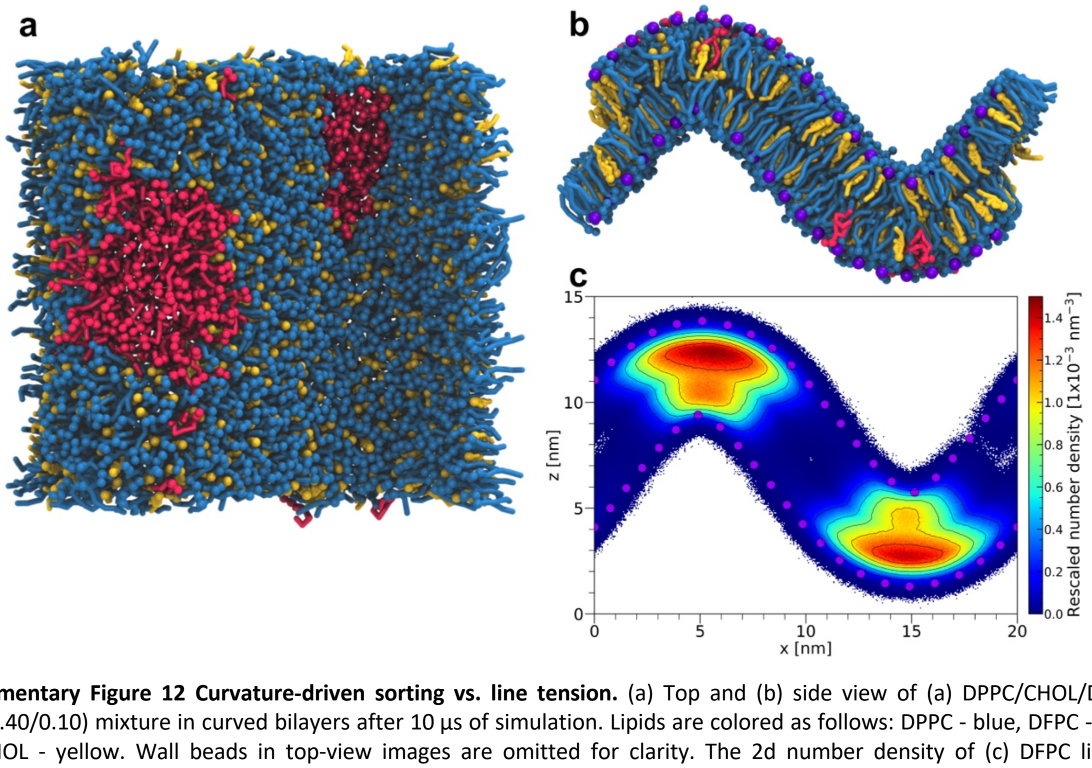


**Supplementary Figure 12 Curvature-driven sorting vs. line tension.** (a) Top and (b) side view of (a) DPPC/CHOL/DFPC (0.50/0.40/0.10) mixture in curved bilayers after 10 µs of simulation. Lipids are colored as follows: DPPC - blue, DFPC - red, and CHOL - yellow. Wall beads in top-view images are omitted for clarity. The 2d number density of (c) DFPC lipids, corresponding to the $L_d$ phase, is computed as an average over the last 2 µs of a single simulation.

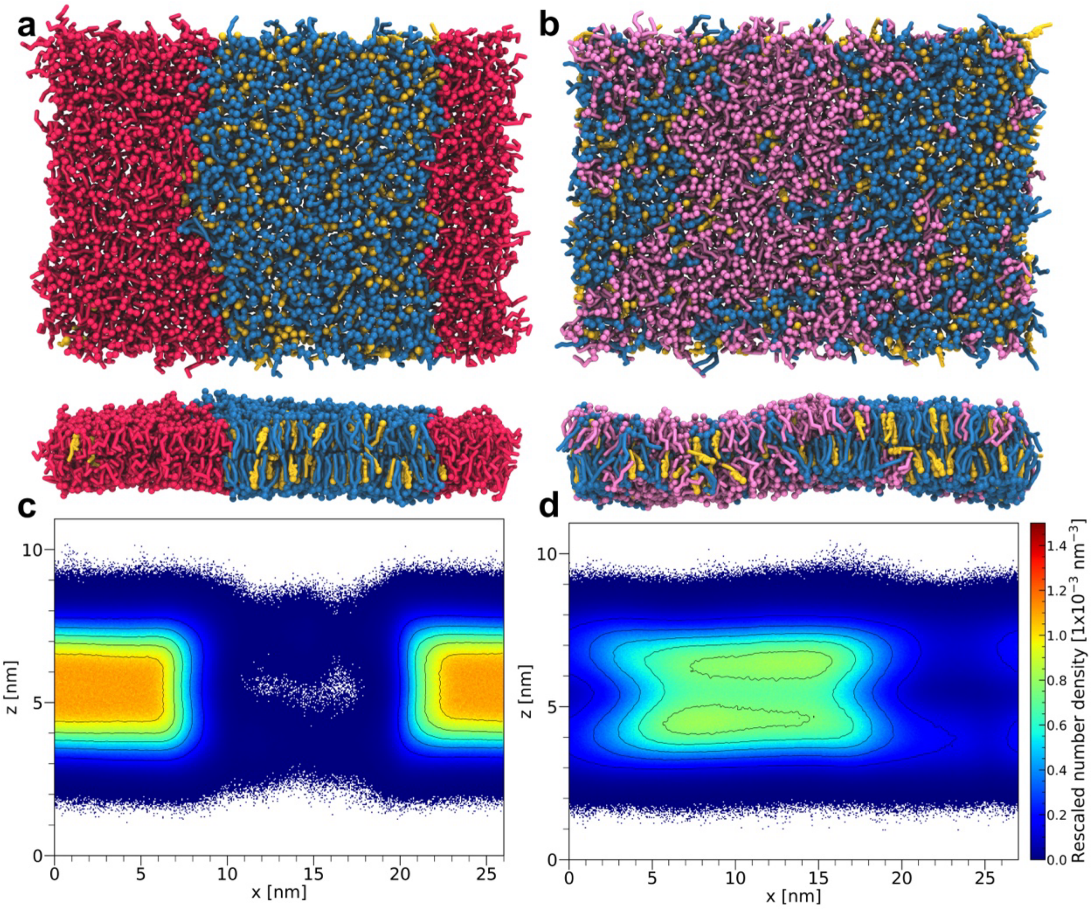


**Supplementary Figure 13 Interleaflet coupling in flat phase separated system.** Top and side view of (a) DPPC/CHOL/DFPC (0.35/0.35/0.30) and (b) DPPC/CHOL/DIPC (0.35/0.35/0.30) mixtures in curved bilayers after 10 µs of simulation. Lipids are colored as follows: DPPC - blue, DFPC - red, CHOL – yellow, and DIPC - pink. The 2d number density of (c) DFPC and (d) DIPC lipids, corresponding to the $L_d$ in each system, is computed as an average over the last 2 µs for a single simulation.

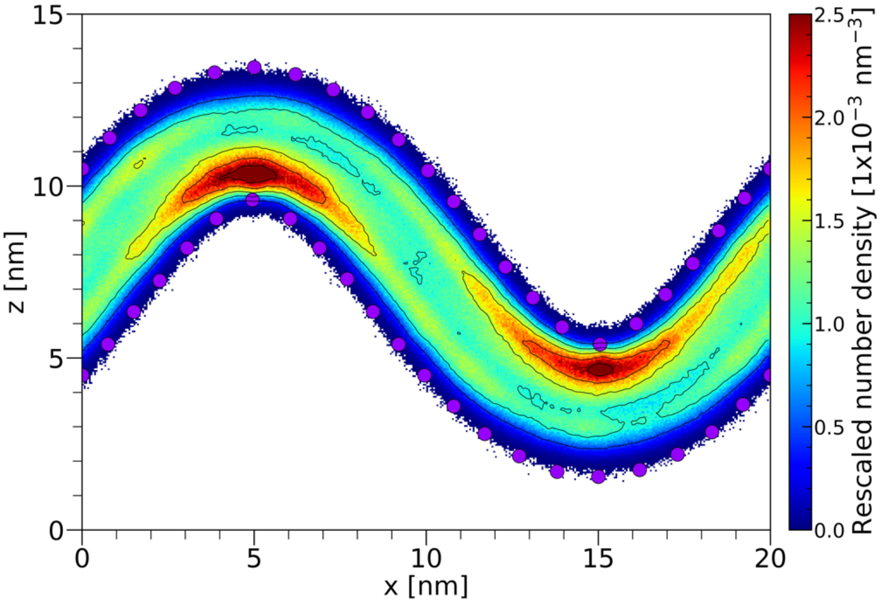


**Supplementary Figure 14 CHOL shows strong enrichment in negative curvature in mixtures with POPC.** The 2D number density of CHOL lipids is computed as an average over the last 2 µs of a single simulation and normalized by the total number of CHOL beads.

### Supplementary Methods

### Simulation setup and parameters

All molecular simulations were performed with the beta release of Martini 3 or final Martini 3 parameters [1] using Gromacs versions 2019.4, 2019.5, 2020.3, 2021.5, and 2022 [2]. Simulation parameters were based on standard values used in the Martini benchmarking by De Jong *et al.* [3].

Curved membrane simulations were set up using a multi-step procedure that ensures minimal influence of the wall beads and simulation parameters on the membrane structure and dynamics. All bilayers were generated using the TS2CG version 1.1.[4] First, to get an estimate for the bilayer thickness, ***flat reference systems*** were generated using an initial box size of 15 x 10 x 10 nm$^3$. After a short vacuum energy minimization using the steepest descent algorithm, all systems were subject to a 50.00 step vacuum NVT equilibration with a 10 fs time step. NVT simulations were performed using the velocity-rescaling thermostat [5] at 310 K with a time constant of 1.0 ps. During equilibration, lipid head groups were fixed to their initial positions to allow proper expansion of lipid tails. Subsequently, all systems were solved and counterions plus 150 mmol NaCl or 50 mmol $CaCl_2$ were inserted using the TS2CG solvation script, reaching a final system size of ≈13.000 CG beads. After another short energy minimization and a 500,000-step NVT equilibration with position restraints, a two-step NPT equilibration with semi-isotropic pressure coupling was performed. During the first round, position restraints on lipids head groups were still applied and the Berendsen barostat [6] was used with a reference pressure of 1 bar and a 10 fs time step for a total of 100,000 steps. All position restraints were released for the second round of NpT equilibration and the time step was increased to 20 fs. The final production run was carried out for at least 500 ns in the NpT ensemble with a 20 fs time step and Parrinello-Rahman barostat [7]. The first 300 ns of the trajectory were excluded from the analysis.

To preserve non-flat membrane shapes, artificial wall beads are introduced, which are designed to only interact with lipid tail beads. Wall beads have almost purely repulsive interaction with C1 and C4h beads using an epsilon of 0.47 and epsilon of 0.1. Due to their artificial nature, wall beads are treated as freeze groups, this means their positions were kept fixed during the entire simulation and are excluded from both temperature and pressure coupling. Simulating ***curved membrane systems*** requires special care because, in the presence of scaffolding wall beads, simulations need to be run in the NpAT ensemble, i.e., with a fixed membrane area, to avoid artifacts at the periodic boundaries. To simulate an approximately tensionless curved bilayer in the NpAT ensemble, the number of lipid molecules needs to be adjusted as the area per lipid changes from a flat to a curved bilayer. First, curved bilayers with an initial box size of 20 x 10 x 15 nm$^3$ were generated using the newly implemented analytic shape function of TS2CG. The bilayer mid-plane is defined by a cosine function with an amplitude of 4 nm and a period of 20 nm. The lipid ratio and membrane thickness are defined based on the previously performed flat reference simulation. Wall beads were placed 0.1 nm above the head group region of each leaflet using a wall bead density of 0.7. Again, all systems were subject to a short vacuum energy minimization and NVT equilibration with applied position restraints on the lipid head groups. After the solvation and insertion of ions, the final system size reaches ≈2,000 CG beads. Subsequently, another short energy minimization and 500,000 steps NVT equilibration with position restraints was performed. In contrast to flat bilayer simulations, a two-step NpAT equilibration is used. Semi-isotropic pressure coupling combined with zero compressibility in the x-y plane realizes a constant membrane area. Again, all position restraints were released during the second round of NpAT equilibration. A short 20 ns production run was performed in the NpT ensemble to verify that the initial number of lipids matches the area of the curved bilayer. The average box size was calculated using *gmx energy* to check for expansion or shrinkage in the x-y dimension, which indicates a mismatch in the number of lipids. If the x dimension changed by more than 0.05%, the initial number of lipids was adjusted and all the previous steps were repeated until the box size remained constant in the NpT ensemble. Finally, 5 µs production runs were carried out in 3 replicas using the NpAT ensemble with Berendsen barostat. Unless mentioned otherwise, the last 2 µs of the 3 replicas were combined and used for analysis.

An additional set of curved membranes with an initial box size of 20 x 20 x 15 $nm^3$ was generated to capture the phase behavior of ternary lipid mixtures. These systems were simulated for 10 µs to allow for proper equilibration of the phases.

To investigate the energy difference of curvature-induced lipid sorting, potentials of mean force (PMFs) for a single lipid we calculated using ***umbrella sampling***. The selected lipid was pulled along the x-dimension of the bilayer with respect to the first wall bead located at x = 0 nm. While the periodic boundary conditions were taken into account, the lipid was free to move in the other two dimensions. 256 umbrella windows were extracted covering the full 20 nm length of the simulation box. Subsequently, each window was simulated for 1 µs with a force constant of 1000 kJ/mol. The same simulation settings as for unbiased curved simulations were used. The PMFs were computed based on the last 800 ns of each window using the weighted histogram analysis method (WHAM) as implemented in *gmx wham* [8] and corrected for the periodic nature of the PMF. Statistical errors were estimated using Bayesian bootstrapping of the complete histogram. Each bootstrapped PMF was defined as zero at x = 0 nm, which roughly corresponds to zero curvature.

### Data analysis

***Area per lipid*** (APL) and ***membrane thickness*** were calculated for flat reference bilayers using the default settings of APL@VORO [9]. For standard lipids, the head group was defined by their phosphate (PO4) beads which were used for the leaflet detection and tessellation. In the case of cardiolipin (CDL), both phosphates (PO41 and PO42) were used. APLs for cardiolipins were then obtained by adding the values for the PO41 and PO42 groups.
To obtain the lipid shape, the ***critical packing parameter*** (CPP) was computed from flat reference bilayers. The CPP is defined as

$$CPP = \frac{v_0}{APL \cdot l_c} \qquad \text{Eq. 1}$$

where $v_0$ is the hydrophobic tail volume and $l_c$ is the tail length. The lipid tail volume $v_0$ was computed for a single lipid using *gmx sasa* with a probe radius of zero and a van-der-Waals radius of 0.264 nm for normal Martini beads. Linker beads were excluded from the volume calculation. The tail length $l_c$ was calculated as an average of both tails (or all four tails for CDL). For a single lipid, the distance in z between the first and every other tail bead was measured. Subsequently, the maximum distance is used to calculate the CPP.

The ***2D densities*** of lipids and ions in curved bilayers were calculated with *gmx densmap* using 400 bins in x- and 300 bins in z-dimension while averaging over the y-direction. The generated data was rescaled by the total number of beads for which the density was calculated.

The ***relative density*** and ***enrichment*** of lipids in curved bilayers were computed to measure the lipid packing and sorting in a curvature gradient. First, a system-specific reference wall with an increased number of wall beads (≈ 52,000) was created with TS2CG, which also generated a file including the mean curvature and area per wall bead. For each leaflet, the number of lipids in closest proximity to each wall bead was calculated. The distance calculation was performed with MDAnalysis [10] considering only lipid head groups, i.e., PO4 beads for standard lipids and GL0 beads for CDLs. Using the area per wall bead *i*, the lipid density per wall bead was calculated (Eq. 2) and normalized by the expected lipid density (Eq. 3) assuming an equal distribution of lipids across each leaflet. Relative densities were computed considering all lipid types present in the bilayer (Eq. 4).

$$density_{@wall_i} = \frac{count_{@wall_i}}{area_{wall_i}} \quad \text{Eq. 2}$$

$$density_{\text{expected}} = \frac{count_{\text{lipids}}}{\sum_i area_{wall_i}} \quad \text{Eq. 3}$$

$$relative\ density_{@wall_i} = \frac{density_{@wall_i}}{density_{\text{expected}}} \quad \text{Eq. 4}$$

Relative enrichment values were obtained by first calculating the relative densities for a single lipid type and then normalizing these densities by the combined relative lipid density of all lipids. The data of all 3 replicas and both leaflets was combined and fitted using a Lowess regression.

Instead of expressing lipid sorting in terms of densities as a function of curvature, the ***sorting index*** was calculated to determine the overall preference for positively or negatively curved membrane regions. As for the relative density, for each leaflet, the closest wall bead for each lipid head group was determined. Depending on the curvature of this wall bead, the lipid resides in negative, zero, or positive curvature. In a mixture of lipids of type *a* and *b*, the ratio of lipids *a* in positively (negatively) curvature membrane regions was calculated by dividing the number of lipids *a* which are closest to wall beads with positive (negative) curvature by the total number of lipids *a* and *b* in the positive (negative) region.

$$ratio_{a_\pm} = \frac{count_{a_\pm}}{count_{a_\pm} + count_{b_\pm}} \quad \text{Eq. 5}$$

The ratio of lipids *a* in negative curvature was subtracted from the ratio of lipids *a* in positive curvature and divided by 2 times the expected (input) ratio of lipid *a*.

$$Sorting\ index_a = \frac{ratio_{a_+} - ratio_{a_-}}{2 \cdot ratio_{a_{expected}}} \quad \text{Eq. 6}$$

A positive sorting index indicates a preference for positively curved membrane regions and vice versa. It should be noted that due to its numeric nature, the sorting index cannot capture simultaneous enrichment of lipids in both positive and negative curvature, which happens in the presence of interleaf let coupling, but will give zero curvature preference.

The ***lipid mixing entropy*** was calculated to measure the disorder-order transition of lipids in mixed bilayers upon curvature-induced sorting. The entropy of mixing was evaluated based on the conditional entropy as introduced by Brandani *et al.* [11] and implemented in Membrainy [12]. As certain lipid types including CDL were missing in the Membrainy library, we used MDAnalysis to calculate the mixing entropy, which also allows for a straightforward leaflet detection. For a membrane containing two or more lipid types, the mixing entropy is defined as

$$S(x_1, .., x_N) = N \sum_{x_i, nb_i} p(x_i, nb_i) \log p(x_i | nb_i) \qquad \text{Eq. 7}$$

where $p(x_i, nb_i)$ is the probability to find a lipid of type $x_i$ neighbored to a lipid of type $nb_i$ and $p(x_i|nb_i)$ describes the conditional probability that a lipid is of type $x_i$ given that its neighbor is of type $nb_i$. For each leaflet, lipid neighborhoods were analyzed by defining a distance vector between lipid head groups (i.e., PO4 beads for standard lipids and GL0 beads for CDLs) of all lipids and the nearest neighboring lipid was determined. The number of nearest neighbors per lipid type is normalized such that the total probability is always 1 and used in Eq. 7. Using the theoretical maximum entropy

$$S_{max} = -\sum p_{x_i} \log p_{x_i} \qquad \text{Eq. 8}$$

where $p_{x_i} \log p_{x_i}$ is the density of a lipid of type $x_i$, the entropy can be normalized, such that $S_{max} = 1$.
Lipid-lipid and lipid-ion ***radial distribution functions*** (RDFs) were calculated using MDAnalysis considering PO4 beads for standard lipids and GL0 for CDLs.
